# The RAB27A effector SYTL5 regulates mitophagy and mitochondrial metabolism

**DOI:** 10.1101/2024.12.30.630740

**Authors:** Ana Lapao, Lauren Sophie Johnson, Laura Trachsel-Moncho, Samuel J. Rodgers, Sakshi Singh, Matthew Yoke Wui Ng, Sigve Nakken, Eeva-Liisa Eskelinen, Anne Simonsen

## Abstract

SYTL5 is a member of the Synaptotagmin-Like (SYTL) protein family that differs from the Synaptotagmin family by having a unique N-terminal Synaptotagmin homology domain that directly interacts with the small GTPase RAB27A. Several SYTL protein family members have been implicated in plasma membrane transport and exocytosis, but the specific function of SYTL5 remains unknown. We here show that SYTL5 is a RAB27A effector and that both proteins localise to mitochondria and vesicles containing mitochondrial material. Mitochondrial recruitment of SYTL5 depends on its interaction with functional RAB27A. We demonstrate that SYTL5-RAB27A positive vesicles containing mitochondrial material, autophagy proteins and LAMP1 form during hypoxia and that depletion of SYTL5 and RAB27A reduces mitophagy under hypoxia mimicking conditions, indicating a role for these proteins in mitophagy. Indeed, we find that SYTL5 interacts with proteins involved in vesicle-mediated transport and cellular response to stress and that its depletion compromises mitochondrial respiration and increases glucose uptake. Intriguingly, SYTL5 expression is significantly reduced in tumours of the adrenal gland, and correlates positively with survival for patients with adrenocortical carcinoma.

## Introduction

The Synaptotagmin-Like (SYTL) protein family consists of 5 members (SYTL1-5), each containing an N-terminal Synaptotagmin homology domain (SHD) and two C-terminal C2 lipid-binding domains. The SHD domains of SYTL1-5 have been found to directly interact with the small GTPase protein RAB27^1,2^. The C2 domain is generally involved in phospholipid binding and interacts with cellular membranes^3^, either in a calcium-dependent or calcium-independent manner^3,4^. Calcium was shown to be required for SYTL3 and SYTL5 phospholipid binding activity^1,5^, whereas SYTL2 binding to phosphatidylserine (PS) was inhibited by calcium^6^.

The SYTL protein family is generally involved in plasma membrane transport and exocytosis^2,4^, mostly through their binding to RAB27 proteins. The main functions of RAB27 are related to vesicle budding, delivery, tethering and fusion with membranes^7^ and its activity is regulated by a cyclic activation and inactivation state depending on it binding to guanosine-5’-triphosphate (GTP) or guanosine diphosphate (GDP), respectively^7^. RAB27A and RAB27B are the two RAB27 isoforms found in vertebrates^7^ and their binding to effector proteins occurs only when they are bound to GTP^7^. SYTL5 binds to the GTP-bound RAB27A form, indicating its possible role in membrane trafficking events^1^.

Functional mitochondria are fundamental for normal cellular metabolism. Mitochondria are the primary source of adenosine triphosphate (ATP) obtained through oxidative phosphorylation (OXPHOS), and are important for cellular calcium homeostasis, lipid metabolism, reactive oxygen species (ROS) generation, and detoxification^8^. Mitochondria can integrate and generate signalling cues to adjust their metabolism and biogenesis to maintain cellular homeostasis^8,9^. Mitochondrial function and health are tightly regulated by several quality control processes involving targeting of parts of mitochondria to lysosomes for degradation, including macromitophagy^10,11^ (selective degradation of mitochondria by autophagy), piecemeal mitophagy^12,13^, mitochondrial-derived vesicles (MDVs)^14,15^, and vesicles derived from the inner mitochondrial membrane (VDIMs)^16^. It has also been found that damaged mitochondria can be directly released to the extracellular space^17^, either contained within extracellular vesicles^18^ or by mitocytosis, where damaged mitochondria are expelled in migrasomes^19^.

Exposure of mitochondria to various stressors can affect their function and lead to severe diseases, such as neurodegenerative disorders and cancer^20,21^. Cancer cells can trigger a change in mitochondrial metabolism to promote cell proliferation and survival by a process known as the Warburg effect, characterised by a switch from OXPHOS to ATP production through glycolysis even under normoxic conditions^22^. Adrenocortical carcinoma (ACC) is a rare type of cancer that develops in the adrenal gland cortex, having a very poor prognosis and limited treatment options^23^. Most patients are diagnosed at advanced cancer stages and present an excess in adrenocortical hormone production^23^. Mitochondria are known to be essential organelles in the synthesis of steroid hormones, including cortisol, since they contain specific enzymes that catalyse steroid synthesis from cholesterol delivered to the mitochondria^24^.

Here we demonstrate that SYTL5 localises to mitochondria in a RAB27A-dependent manner and that both proteins regulate mitophagy and cellular metabolism. Cells lacking SYTL5 undergo a shift from mitochondrial oxygen consumption to glycolysis, which may explain the correlation between low SYTL5 expression and poor survival of ACC patients.

## Results

### SYTL5 localises to mitochondria and endolysosomal compartments

SYTL5 was identified as a putative candidate in a screen for lipid-binding proteins involved in the regulation of mitophagy^25^. To characterise the cellular localisation and function of SYTL5, we generated U2OS cells with stable inducible expression of SYTL5-EGFP as there are no antibodies recognising endogenous SYTL5 and our efforts to tag endogenous SYTL5 using CRISPR/Cas9 were also unsuccessful. Intriguingly, live cell imaging analysis revealed that SYTL5-EGFP co-localised with filamentous structures positive for MitoTracker red **(Figure 1A)**, which labels active mitochondria. Moreover, several SYTL5-EGFP positive vesicles were observed, including some small and highly mobile SYTL5-EGFP vesicles moving along the mitochondrial network **(Supplementary video 1 and arrows in Figure 1B)**. SYTL5-EGFP staining at the plasma membrane was also seen, particularly in cells with higher expression levels **(Figure 1C)**, which may suggest a role for SYTL5 in secretion to the plasma membrane.

**Figure 1.**
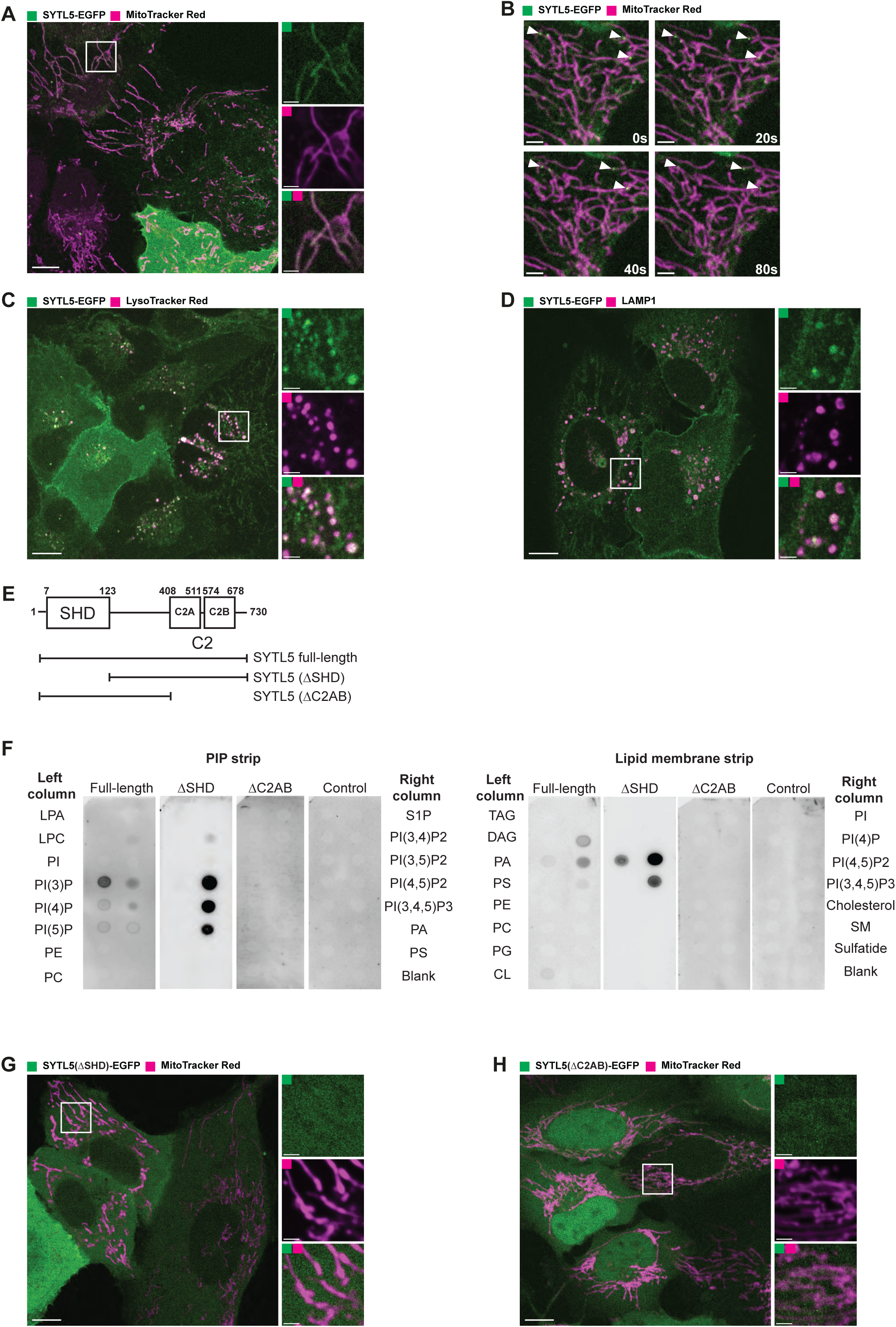
Lipid binding protein SYTL5 localises to mitochondria and endo-lysosomes. A. Live confocal microscopy imaging of U2OS stably expressing SYTL5-EGFP co-stained with 50 nM MitoTracker red. SYTL5-EGFP expression was induced for 24 h using 100 ng/ml doxycycline. MitoTracker red was added 30 min before imaging. Scale bars: 10 μm, 2 μm (insets). B. Time-lapse video frames (Supplementary video 1) tracking SYTL5 vesicle movement along filaments positive for MitoTracker red (arrows). SYTL5-EGFP expression was induced for 24 h using 100 ng/ml doxycycline. Scale bars: 3 μm. C. Live confocal microscopy imaging of U2OS stably expressing SYTL5-EGFP co-stained with 50 nM LysoTracker red. SYTL5-EGFP expression was induced for 24 h using 100 ng/ml doxycycline. LysoTracker red was added 30 min before imaging. Scale bars: 10 μm, 2 μm (insets). D. Confocal imaging of U2OS stably expressing SYTL5-EGFP stained for endogenous LAMP1. Nuclei were stained using Hoechst. Scale bars: 10 μm, 2 μm (insets). E. Overview of domain structure of SYTL5 full length, ΔSHD and ΔC2AB mutants. F. 3xFLAG fusion proteins expressed in U2OS cells (SYTL5-EGFP-3xFLAG, SYTL5 (ΔSHD)-EGFP-3xFLAG and SYTL5 (ΔC2AB)-EGFP-3xFLAG) were immunoprecipitated and added to membranes spotted with lipids found in cell membranes: LPA (lipoprotein A); LPC (lysophosphatidylcholine); PI (phosphatidylinositol); PI(3)P (phosphatidylinositol 3-phosphate); PI(4)P (phosphatidylinositol 4-phosphate); PI(5)P (Phosphatidylinositol 5-phosphate); PE (phosphatidylethanolamine); PC (phosphatidylcholine); S1P (sphingosine-1-phosphate); PI(3,4)P2 (phosphatidylinositol 3,4-bisphosphate); PI(3,5)P2 (phosphatidylinositol 3,5-bisphosphate); PI(4,5)P2 (phosphatidylinositol 4,5-bisphosphate); PI(3,4,5)P3 (phosphatidylinositol 3,4,5 –triphosphate); PA (phosphatidic acid); PS (Phosphatidylserine); TAG (triglyceride); DAG (diglyceride); PG (phosphatidylglycerol); CL (cardiolipin); Cholesterol; SM (sphingomyelin); Sulfatide. G. Live confocal microscopy imaging of U2OS stably expressing SYTL5 (ΔSHD)-EGFP-3xFLAG co-stained with 50 nM MitoTracker red added 30 min before imaging. Scale bars: 10 μm, 2 μm (insets). H. Live confocal microscopy imaging of U2OS stably expressing SYTL5 (ΔC2AB)-EGFP-3xFLAG co-stained with 50 nM MitoTracker red added 30 min before imaging. Scale bars: 10 μm, 2 μm (insets).

To determine the identity of the SYTL5 positive vesicles, U2OS SYTL5-EGFP cells were infected with a panel of lentiviral RAB GTPase constructs representing different cellular compartments **(Supplementary Figure 1A-F)** and analysed by live cell imaging. SYTL5-EGFP positive structures were observed to co-localise with vesicles positive for RAB4 and RAB11, two GTPases that mainly localise to recycling endosomes^26^ and to some extent with RAB5, mostly localised to early endosomes^26^ **(Supplementary Figure 1A-C)**. Co-localisation was also observed with RAB7, a marker of late endosomes, lysosomes and autophagosomes^26^ and partially with RAB9, which associates with late endosomes and mediates the transport between late endosomes and the *trans*-Golgi network^26^ **(Supplementary Figure 1D-E)**. SYTL5-EGFP showed little or no co-localisation with RAB6 positive structures, representing Golgi-derived vesicles **(Supplementary Figure 1F)**. In line with an endocytic identity of SYTL5-EGFP positive vesicles, they were also positive for lysosomal markers, including LysoTracker red, which labels acidic compartments in the cell **(Figure 1C)**, and the lysosomal membrane marker LAMP1 **(Figure 1D)**.

Thus, based on live cell imaging analysis we conclude that SYTL5 localises to the mitochondrial network and to mitochondria-associated vesicles that partly overlap with endolysosomal compartments.

### Mitochondrial localisation of SYTL5 requires both the RAB27-binding SHD domain and the lipid-binding C2 domains

To investigate whether the observed intracellular localisation of SYTL5-EGFP depends on its binding to RAB27 and/or lipids, we generated U2OS cells with stable expression of EGFP-3xFLAG-tagged wild type SYTL5 or SYTL5 lacking either the RAB27-binding SHD domain (SYTL5(ΔSHD)) or the two lipid-binding C2 domains (SYTL5(ΔC2AB)) **(Figure 1E)**.

To validate the SYTL5(ΔC2AB) mutant and determine the lipid-binding specificity of SYTL5, lysates from these cells were incubated with membranes containing various phosphoinositides and other lipids. Full-length SYTL5 was observed to bind to mono-phosphorylated phosphoinositides (PI(3)P, PI(4)P, PI(5)P), as well as PI(4,5)P2, PI(3,4,5)P3 and to some extent to phosphatidic acid (PA) and cardiolipin (CL) **(Figure 1F)**. As expected, upon removal of the C2 domains, specific binding to all lipid species was lost, while SYTL5 lacking the SHD domain retained the ability to bind to lipids **(Figure 1F)**.

The SYTL5(ΔSHD)-EGFP and SYTL5(ΔC2AB)-EGFP expressing cell lines were then incubated with MitoTracker red and analysed by live cell imaging. Indeed, the SYTL5 colocalisation to mitochondria was lost and both SYTL5(ΔSHD)-EGFP and SYTL5(ΔC2AB)-EGFP were dispersed in the cytosol **(Figure 1G-H),** indicating that both the RAB27-interacting SHD domain and the lipid binding C2 domains are required for mitochondrial localisation of SYTL5.

### Mitochondrial localisation of SYTL5 requires RAB27A GTPase activity

Given that the SHD domain of SYTL5 has been shown to bind RAB27A^1^ and our observations that the SHD domain is required for mitochondrial localisation of SYTL5, we asked whether RAB27A activity is required for SYTL5 recruitment to mitochondria. U2OS SYTL5-EGFP cells were infected with a mScarlet-RAB27A lentiviral construct to generate a stable cell line. Co-expression of SYTL5-EGFP and mScarlet-RAB27A resulted in a clear co-localisation of both proteins to filaments positive for MitoTracker DeepRed (DR) **(Figure 2A)**. The mitochondrial localisation of SYTL5-EGFP and mScarlet-RAB27A was confirmed by correlative light and EM (CLEM) analysis using the same cell line (**Figure 2B, zoom in 1)**. Intriguingly, the mitochondrial localisation of SYTL5-EGFP was enhanced when co-expressed with mScarlet-RAB27A compared to cells expressing SYTL5-EGFP only **(Figure 1A-B)**, indicating that RAB27A might facilitate mitochondrial recruitment of SYTL5. Indeed, in U2OS cells with stable expression of mScarlet-RAB27A only, RAB27A strongly co-localised with MitoTracker green **(Supplementary Figure 2A)**, demonstrating its mitochondrial targeting. Moreover, SYTL5-EGFP and mScarlet-RAB27A were both detected in the mitochondrial fraction (containing TIM23 and COXIV) of cells expressing SYTL5-EGFP or mScarlet-RAB27A **(Supplementary Figure 2B)**. Importantly, the mitochondrial localisation of SYTL5 and RAB27A is not specific to U2OS cells as HeLa cells with stable expression of SYTL5-EGFP or mScarlet-RAB27A show a similar expression pattern to U2OS cells, as demonstrated by co-localisation of SYTL5-EGFP with MitoTracker Red and plasma membrane staining in cells having higher expression **(Supplementary Figure 2C)** and of mScarlet-RAB27A with MitoTracker Green **(Supplementary Figure 2D).** All further experiments were conducted in U2OS cells.

**Figure 2.**
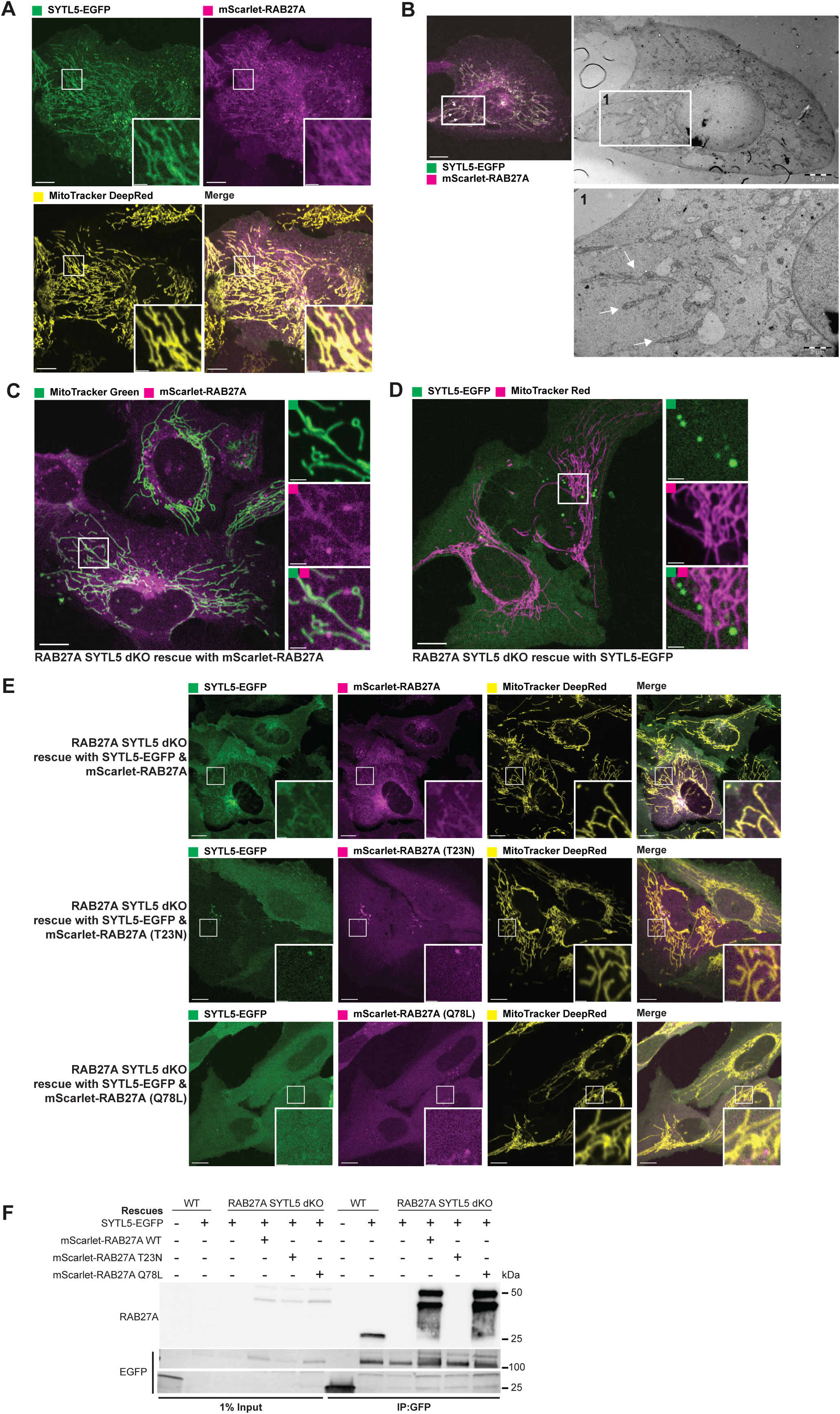
Mitochondrial localisation of SYTL5 requires RAB27A GTPase activity. A. SYTL5-EGFP expression was induced for 24 h with 100 ng/ml doxycycline in U2OS cells with stable inducible expression of SYTL5-EGFP and constitutive expression of mScarlet-RAB27A. Cells were co-stained with MitoTracker DR for 30 min before live imaging. Arrows indicate mitochondrion filaments. Scale bars: 10 μm, 2 μm (insets). B. CLEM analysis of the cells described in A. Before imaging, cells growing in monolayer were fixed in warm (≈37°C) 3.7% paraformaldehyde in 0.2 M HEPES (pH 7). After fixation, cells were imaged using a confocal microscope to acquire Z-stack of optical sections and DIC images to locate the cells of interest. The cells were finally fixed using 2% glutaraldehyde in 0.2 M HEPES (pH 7.4) for 120 min before sample preparation for TEM. Arrows in panel 1 indicate mitochondrion filaments. Scale bars: 15 μm (left), 5 μm (middle) and 2 μm (right). C. U2OS dKO cells were rescued with mScarlet-RAB27A and co-stained with 50 nM MitoTracker green for 30 min before imaging. Scale bars: 10 μm, 2 μm (insets). D. U2OS dKO cells were rescued with SYTL5-EGFP and co-stained with 50 nM MitoTracker red for 30 min before imaging. Scale bars: 10 μm, 2 μm (insets). E. Live confocal microscopy imaging of U2OS dKO cells rescued with SYTL5-EGFP and mScarlet-RAB27A (upper panel); SYTL5-EGFP and mScarlet-RAB27A-T23N (middle panel) or SYTL5-EGFP and mScarlet-RAB27A-Q78L (lower panel). All cells were co-stained with 50 nM MitoTracker DR for 30 min before imaging. Scale bars: 10 μm, 2 μm (insets). F. Lysates from U2OS cells stably expressing EGFP (control) and U2OS dKO cells expressing SYTL5-EGFP and/or mScarlet-RAB27A (wild-type, T23N or Q78L mutants) were immunoprecipitated using GFP-Trap beads and analysed by western blot using RAB27A and EGFP antibodies.

To further characterise mitochondrial recruitment of SYTL5 and RAB27A and elucidate their role at the mitochondrion, we used CRISPR/Cas9 to knock out (KO) RAB27A and SYTL5 in U2OS cells **(Supplementary Figure 3A-D)**. This double-KO (dKO) cell line was further transduced with lentiviral constructs to constitutively express SYTL5-EGFP, mScarlet-RAB27A, or both, and their mitochondrial localisation was analysed by live cell imaging **(Figure 2C-E)**. Intriguingly, mScarlet-RAB27A co-localised extensively with MitoTracker green and to small vesicles dispersed in the cytoplasm and located at or near mitochondria **(Figure 2C)**, indicating that SYTL5 is not required for RAB27A mitochondrial localisation. In contrast, SYTL5-EGFP was not recruited to mitochondria when expressed alone in dKO cells, although some SYTL5-EGFP vesicles were seen near mitochondrial structures **(Figure 2D)**. Upon co-expression of SYTL5-EGFP and mScarlet-RAB27A in dKO cells the mitochondrial localisation of SYTL5-EGFP was rescued **(Figure 2E),** demonstrating that mitochondrial recruitment of SYTL5 is dependent on RAB27A.

Since SYTL5 has previously been described as a RAB27A effector^1^, we asked whether its mitochondrial recruitment depends on binding to the GTP-bound form of RAB27A. To address this, dKO cells expressing SYTL5-EGFP were rescued with the constitutively active (RAB27A-Q78L) or inactive (RAB27A-T23N) mutant forms of mScarlet-RAB27A. As expected, SYTL5-EGFP interacted specifically with RAB27A wild type (WT) and RAB27A-Q78L, as well as with endogenous RAB27A in control cells, while no interaction was detected with RAB27A-T23N, as assessed by GFP-pulldown **(Figure 2F)**. To our surprise, live cell imaging of the same cells revealed that, in contrast to mScarlet-RAB27A WT, neither the RAB27A Q78L nor the T23N mutant localised to the mitochondrial network when expressed together with SYTL5-EGFP in dKO cells **(Figure 2E)**. Also, SYTL5-EGFP failed to localise to mitochondria in cells expressing RAB27A Q78L, further demonstrating that mitochondrial recruitment of SYTL5 depends on mitochondrial RAB27A localisation **(Figure 2E).** Taken together, our data indicate that the GTPase activity of RAB27A is required for its mitochondrial localisation, which facilitates further recruitment of SYTL5.

### SYTL5 interacts with proteins involved in vesicle-mediated transport and cellular response to stress

As few molecular interactors of SYTL5 are known^1^, we analysed the SYTL5 interactome by immunoprecipitating SYTL5-EGFP or an EGFP control, followed by mass spectrometry (MS) and protein-hit enrichment analysis. RAB27A was one of the most significant hits found, providing evidence that the IP was successful, together with several proteins known to participate in secretion such as PDCD6IP^27^ **(Figure 3A)**. In total, 163 proteins were identified as significant SYTL5-EGFP interactors compared to the EGFP control **(Figure 3A**, **Table 1)** and were subjected to Gene Ontology (GO) term enrichment analysis. Considering GO enrichment within the subcellular compartment ontology (GO-CC), we discovered the following compartments as enriched (**Figure 3B**) (number of hits for each category in parentheses): cytoskeleton (43), intracellular vesicle (38), extracellular vesicle (36), plasma membrane-bound cell projection (33), endosome (18) and focal adhesion (17). Furthermore, for biological processes, encoded in the GO-BP subontology, we discovered the following categories as enriched **(Figure 3A and C):** vesicle-mediated transport (38), cellular response to stress (34), response to oxygen-containing compound (31), intracellular protein transport (22), secretion (21), autophagy (13), stress-activated MAPK cascade (10), cellular response to oxidative stress (9), reactive oxygen species metabolic process (7), regulation of mitochondrion organization (6), endosome organization (4) and protein insertion into mitochondrial membrane (3). Taken together, the SYTL5 interactome implies a role for SYTL5 in mitochondrial processes and cellular response to stress, in line with its localisation to mitochondria, as well as a possible role in secretion and endocytosis, in line with its localisation to endocytic compartments and the plasma membrane **(Figure 1)**.

**Figure 3.**
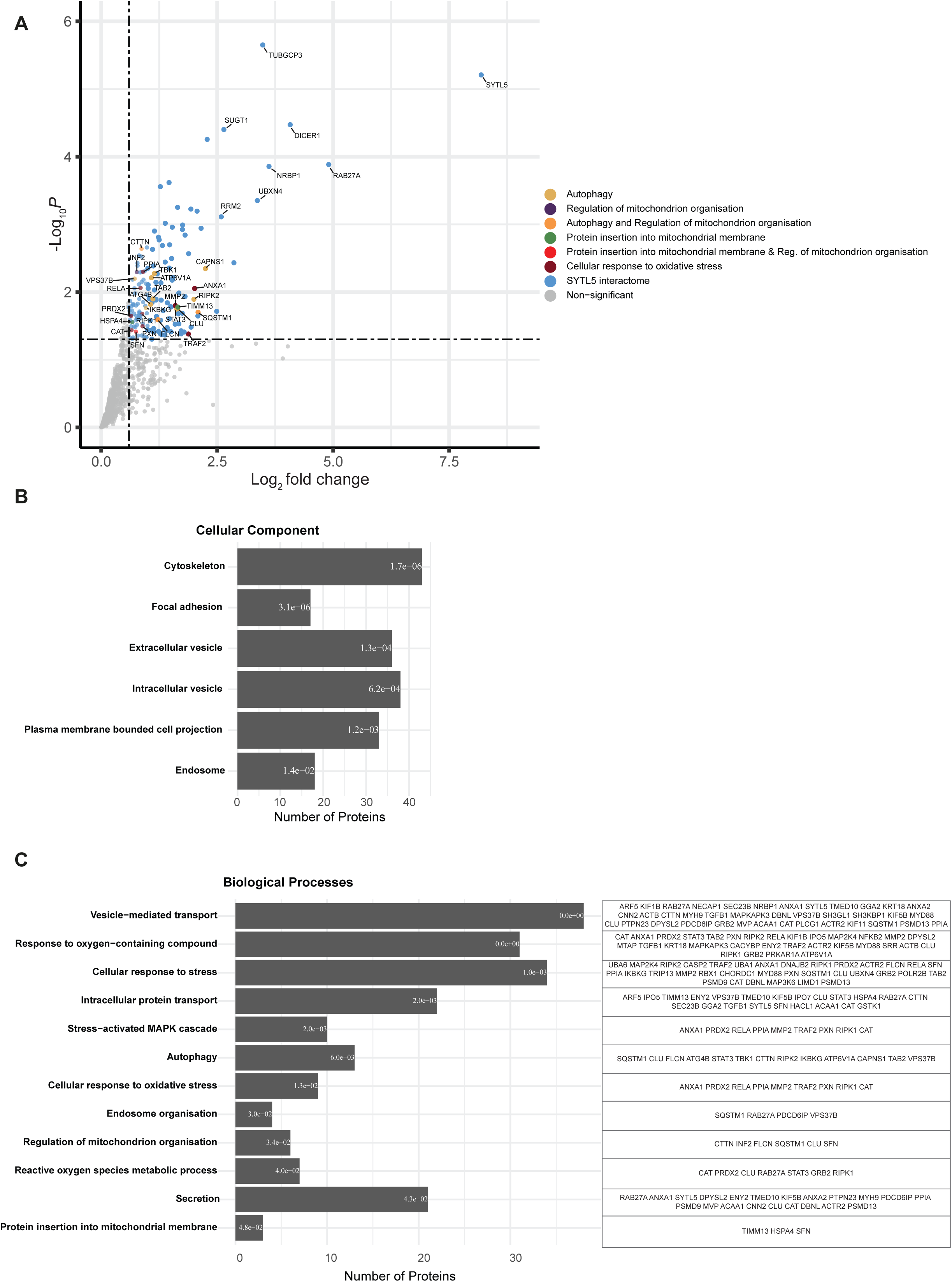
SYTL5 interacts with proteins involved in vesicle-mediated transport and cellular response to stress. A. SYTL5 interactome. SYTL5-EGFP expression in U2OS cells was induced for 24 h using 100 ng/ml doxycycline, followed by immunoprecipitation of SYTL5-EGFP using GFP-Trap beads and identification of co-purified proteins by MS analysis. The list of co-purified proteins was compared with that obtained from cells expressing EGFP and only significant (p<0.05) SYTL5-EGFP specific protein hits are highlighted in colour. Protein hits falling into at least one of the biological processes; autophagy, regulation of mitochondrion organisation, protein insertion into mitochondrial membrane or cellular response to oxidative stress, are colour coded according to plot legend. All other significant hits are indicated in blue. Non-significant (p>0.05) are indicated in grey. Data are from 3 biological replicates. B. GO cellular compartment term enrichment of significant protein hits co-purified with SYTL5-EGFP. Corresponding enrichment false discovery rate (FDR) value is represented inside each bar and bars are ordered from smallest to largest FDR (q-value) from top to bottom. C. GO biological processes term enrichment of significant protein hits co-purified with SYTL5-EGFP.Corresponding enrichment FDR value is represented inside each bar and bars are ordered from smallest to largest FDR (q-value) from top to bottom. Proteins corresponding to each biological process category are listed in the table to the right.

**Table 1.**
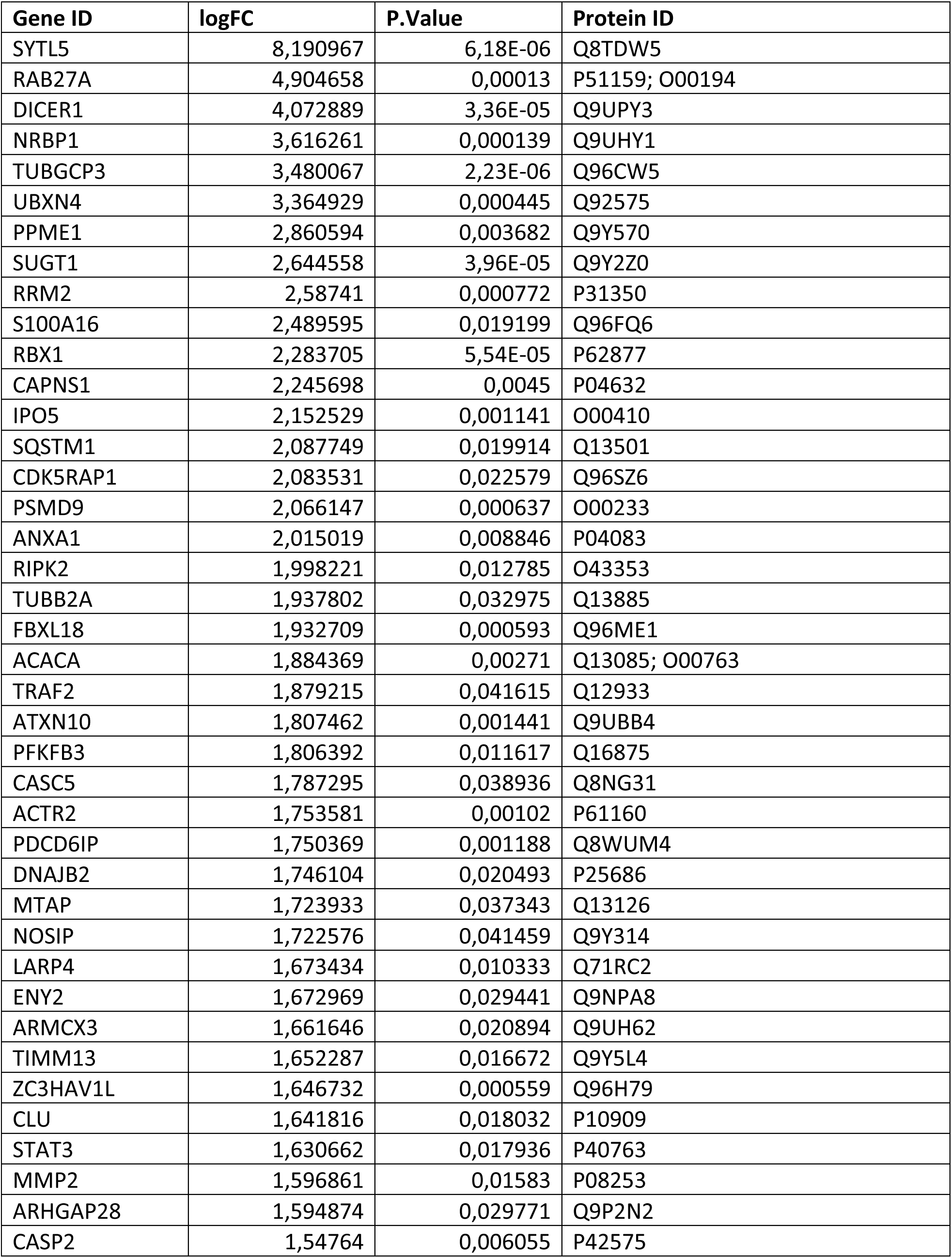
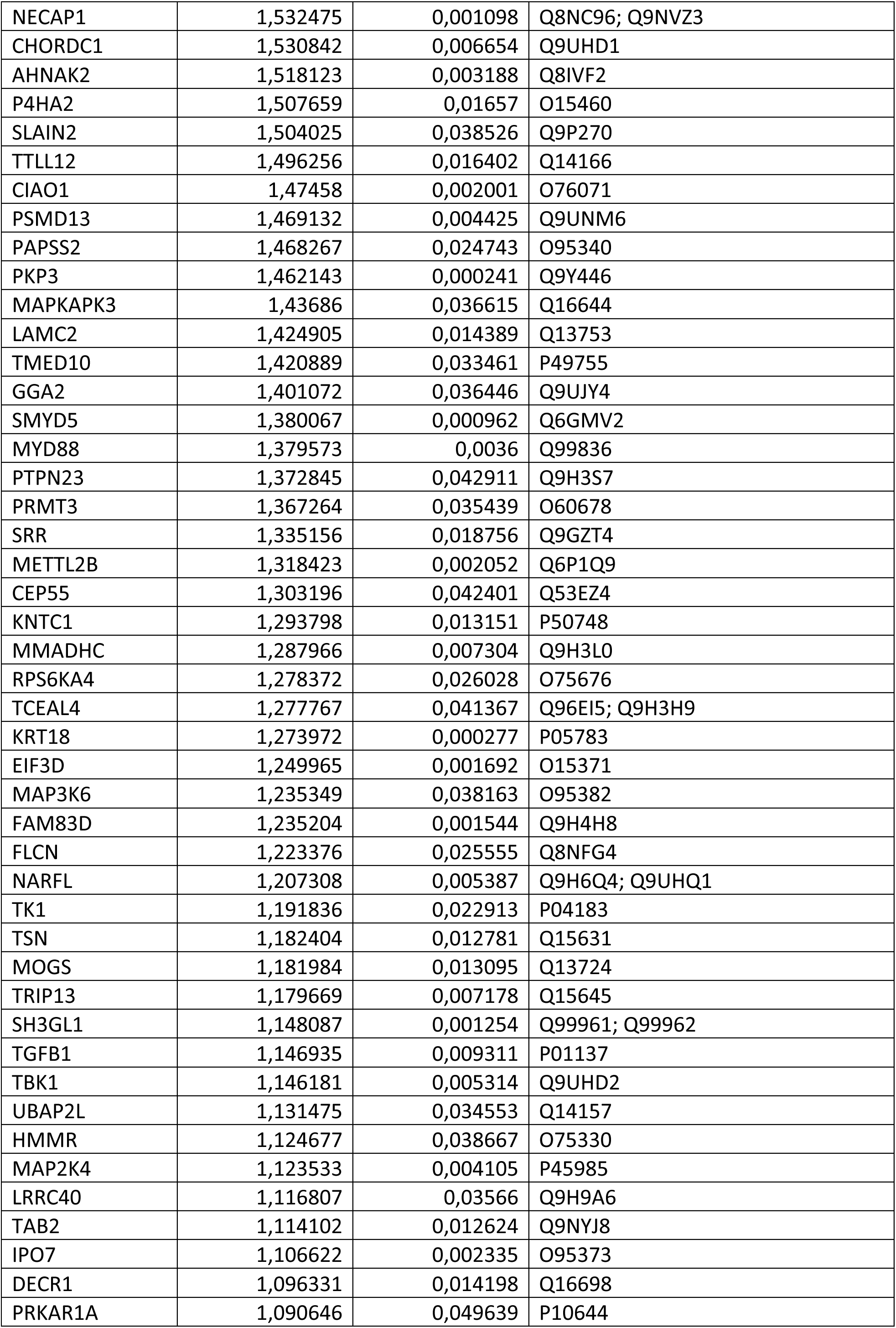
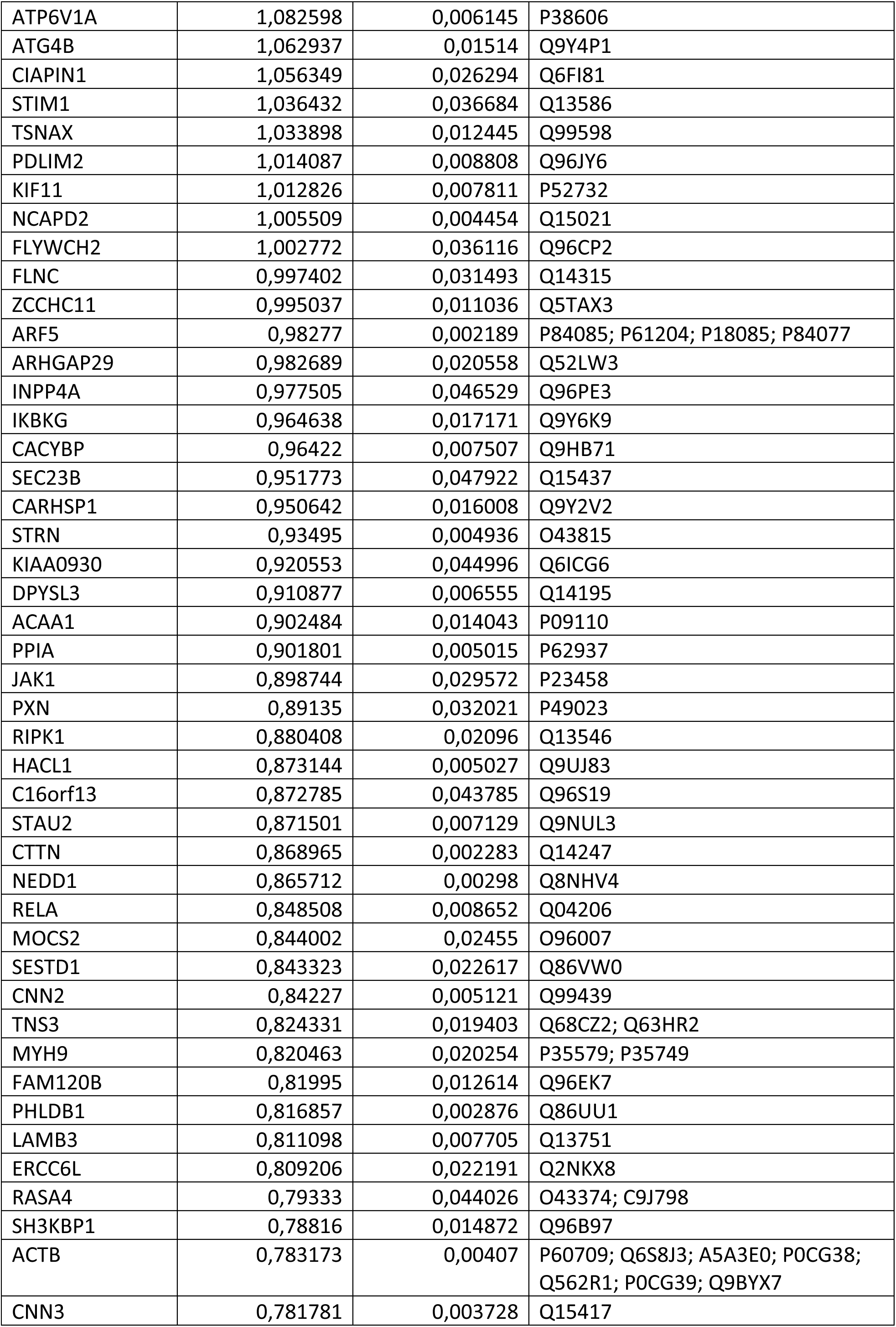
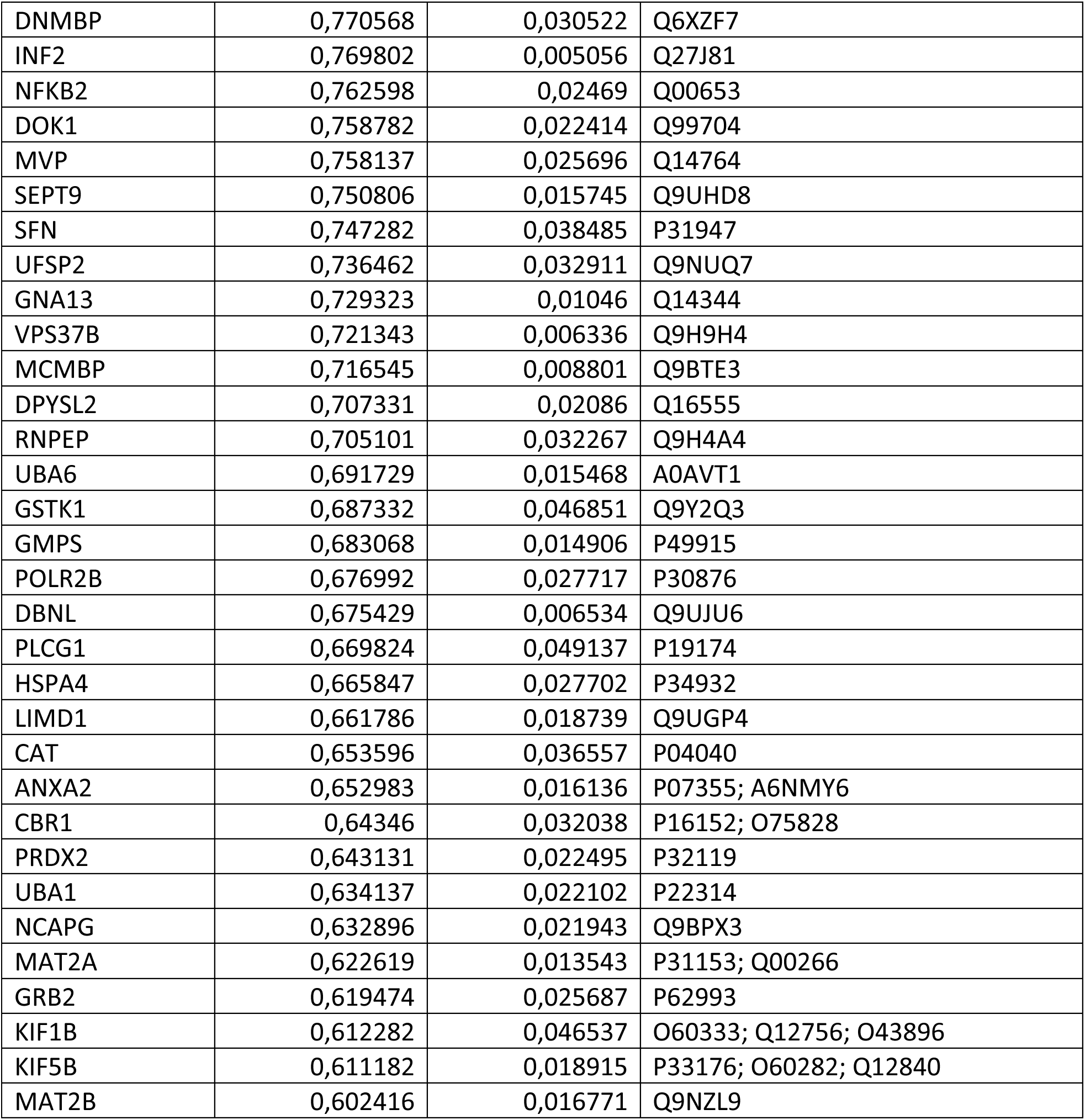
SYTL5 interactome. Proteins co-immunoprecipitated with SYTL5-EGFP vs EGFP expressing control.

### SYTL5-RAB27A positive vesicles contain mitochondrial components

Several proteins involved in autophagy were identified as specific SYTL5 interactors, including SQSTM1/p62, ATG4B, TBK1, and ATP6V1A **(Figure 3A)**, and given the mitochondrial localisation of SYTL5, we first investigated a possible role of SYTL5 in mitophagy. To induce mitophagy, dKO cells expressing mScarlet-RAB27A and SYTL5-EGFP were subjected to hypoxia or the drugs DFP (deferiprone, an iron chelator) or DMOG (Dimethyloxalylglycine, a prolyl-4-hydroxylase inhibitor) that mimics hypoxia by activation of HIF1α. We noticed an increase in the number and size of vesicles positive for SYTL5-EGFP and mScarlet-RAB27A that also contained MitoTracker Deep Red (MitoTracker DR) in all conditions, compared to control cells **(Figure 4A).**

**Figure 4.**
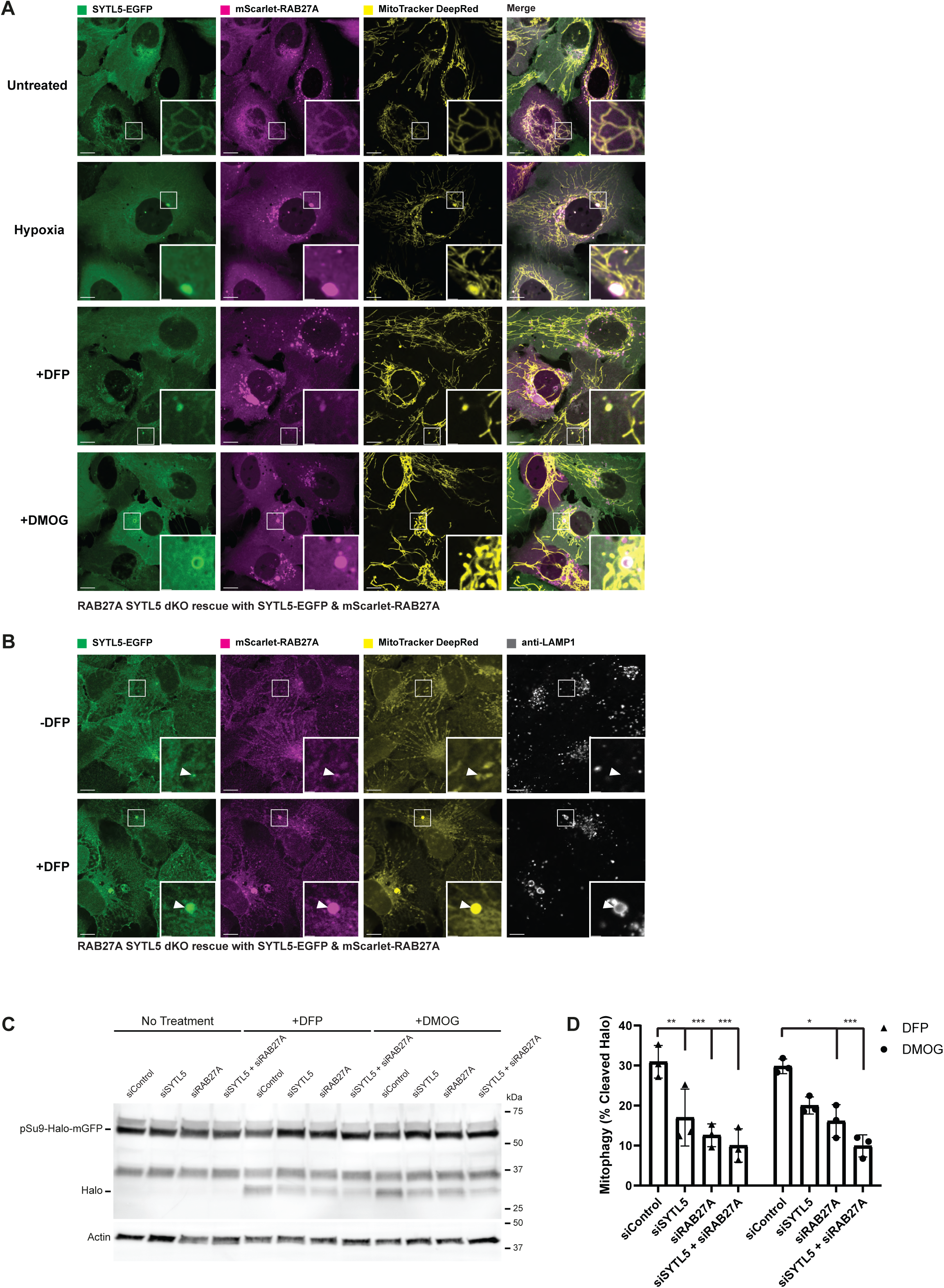
Mitochondrial localisation of SYTL5 and RAB27A is not perturbed by autophagy inhibition or mitophagy stimulation. A. SYTL5/RAB27A dKO U2OS cells rescued with SYTL5-EGFP and mScarlet-RAB27A were untreated (upper panel) or treated with 1 mM DFP (2^nd^ panel), hypoxia (1% O_2_) (3^rd^ panel) or 1 mM DMOG (4^th^ panel) for 24 h. All cells were stained with 50 nM MitoTracker DR 30 min prior to live confocal microscopy imaging. Scale bars: 10 μm, 2 μm (insets). B. SYTL5/RAB27A dKO U2OS cells rescued with SYTL5-EGFP and mScarlet-RAB27A were treated or not with 1 mM DFP for 24 h. Cells were stained with 50 nM MitoTracker DR prior to fixation. Fixed cells were stained with a LAMP1 antibody and imaged by confocal microscopy. Scale bars: 10 μm, 2 μm (insets). Arrowheads point to SYTL5/RAB27A-positive structures. C. U2OS cells expressing pSu9-Halo-mGFP were transfected with 20 nM control siRNA, SYTL5 siRNA #1 or RAB27A siRNA #1 for 48 h before 20 min incubation with the TMR-conjugated Halo ligand followed by 24 h incubation with 1 mM DFP or 1 mM DMOG. Cell lysates were analysed by western blot for the Halo tag. Actin was used as a loading control. D. Quantification of data in C. Error bars represent the mean with standard deviation between replicates (n=3). Significance was determined by two-way ANOVA followed by Tukey’s multiple comparison test. * = p< 0.05, ** =p< 0.01, *** = p<0.001.

HIF-1α is a transcription factor that is a main driver of the Warburg effect^15^ and an inducer of mitophagy via upregulation of the outer mitochondrial membrane proteins BNIP3 and BNIP3L, which bind to LC3. To identify the nature of the SYTL5/RAB27A/MitoTracker DR positive vesicular structures that increase with HIF-1α stabilisation, dKO cells expressing mScarlet-RAB27A and SYTL5-EGFP were immunostained with antibodies against various autophagy markers. Intriguingly, while observing strong co-localisation of SYTL5/RAB27A/MitoTracker DR positive vesicles with LAMP1 (**Figure 4B**), the same structures did not co-localise with LC3B or the autophagy receptor p62 (**Supplementary Figure 4A**). However, as both LC3 and p62 are degraded in the lysosome, we added the lysosomal V-ATPase inhibitor Bafilomycin A1 (BafA1) for the last 4 hrs of DFP treatment, resulting in the presence of some LC3 and p62 positive SYTL5/RAB27A/MitoTracker DR vesicles (**Supplementary Figure 4B**). It should be noted that the SYTL5/RAB27A/MitoTracker-positive vesicle structures do not always colocalise with LAMP1, LC3 and p62, and that the phenotype of these cells is varied. The mitochondrial localisation of SYTL5 and RAB27A was not affected in cells treated with the ULK1 inhibitor MRT68291 or with the class III phosphoinositide 3-kinase VPS34 inhibitor IN1 (**Supplementary Figure 4C**), indicating that SYTL5 and RAB27A are recruited to mitochondria independent of active autophagy. Thus, our data show that SYTL5 and RAB27A-positive vesicles containing mitochondrial material and autophagic proteins form upon HIF-1α induced mitophagy.

### SYTL5 and RAB27A function as positive regulators of selective mitophagy

To investigate whether SYTL5 and RAB27A are required for the turnover of mitochondrial proteins in hypoxic conditions, we took advantage of two mitophagy reporter U2OS cell lines, one expressing the mitochondrial matrix reporter, pSu9-Halo-mGFP^28^ and one expressing the mitochondrial targeting signal of the matrix protein NIPSNAP1 tagged with EGFP-mCherry^29^ (hereafter referred to as IMLS cells).

The pSu9-Halo-mGFP cells allow a measure of mitophagy flux via the production of a cleaved Halo Tag band, which is stable in lysosomes when bound to the ligand (Halo^TMR^). Activation of HIF1α by treatment with DFP or DMOG resulted in the appearance of cleaved Halo^TMR^ **(Figure 4C-D)** consistent with activation of mitophagy. The use of small interfering RNA (siRNA) to deplete SYTL5, RAB27A or both simultaneously in these cells, resulted in a significant reduction in the amount of free Halo^TMR^ **(Figure 4C-D)** highlighting a reduction in mitophagy. Depletion of SYTL5 and/or RAB27A did not affect the DFP or DMOG-induced HIF1α activation, as analysed by BNIP3L expression levels **(Supplementary Figure 5A).** Thus, both SYTL5 and RAB27A are required for the lysosomal turnover of mitochondrial components in response to HIF1α activation and not for the HIF1α mediated induction of mitophagy.

In contrast, we did not observe a significant difference in mitophagy levels in IMLS cells depleted of SYTL5 or RAB27A treated with DFP to induce Parkin-independent mitophagy, or in IMLS cells with stable expression of Parkin treated with the uncoupler carbonyl cyanide m-chlorophenyl hydrazone (CCCP) to induce Parkin-dependent mitophagy **(Supplementary Figure 5B-E)**, together indicating that macro-mitophagy proceeds independently of SYTL5 and RAB27A. In line with this, the starvation-induced autophagic flux was not affected in cells lacking SYTL5 **(Supplementary Figure 5F)**.

Taken together, our results indicate that SYTL5 and RAB27A function to regulate the lysosomal turnover of selected mitochondrial proteins in response to hypoxic conditions.

### SYTL5 regulates mitochondrial respiration

Our finding of mitochondrial material within SYTL5-RAB27A positive structures upon induction of mitochondrial stress through DFP and hypoxia **(Figure 4A)**, together with reduced turnover of the Fo-ATPase subunit 9 upon knockdown of SYTL5 and RAB27A **(Figure 4C-D)** and the identification of several proteins involved in the cellular response to stress **(Figure 3C)**, including mitochondrial proteins (e.g., DECR1 and TIMM13), in the SYTL5 interactome **(Figure 3A)** led us to investigate a potential role of SYTL5 and RAB27A in mitochondrial function and bioenergetics.

The cellular oxygen consumption rate (OCR) can be measured with the Seahorse analyser upon sequential addition of different inhibitors (oligomycin, CCCP, antimycin and rotenone) that block the function of specific electron transport chain (ETC) complexes^30^. SYTL5 KO cells had a significant decrease in both basal and ATP-linked respiration compared to WT U2OS cells **(Figure 5A-B),** indicating reduced mitochondrial OXPHOS activity. As for RAB27A, both basal respiration and ATP-linked OCR were significantly reduced in cells depleted of RAB27A (RAB27A KO and RAB27A/SYTL5 dKO) relative to the control **(Figure 5A-B),** indicating an important function of these proteins in regulation of mitochondrial respiration.

**Figure 5.**
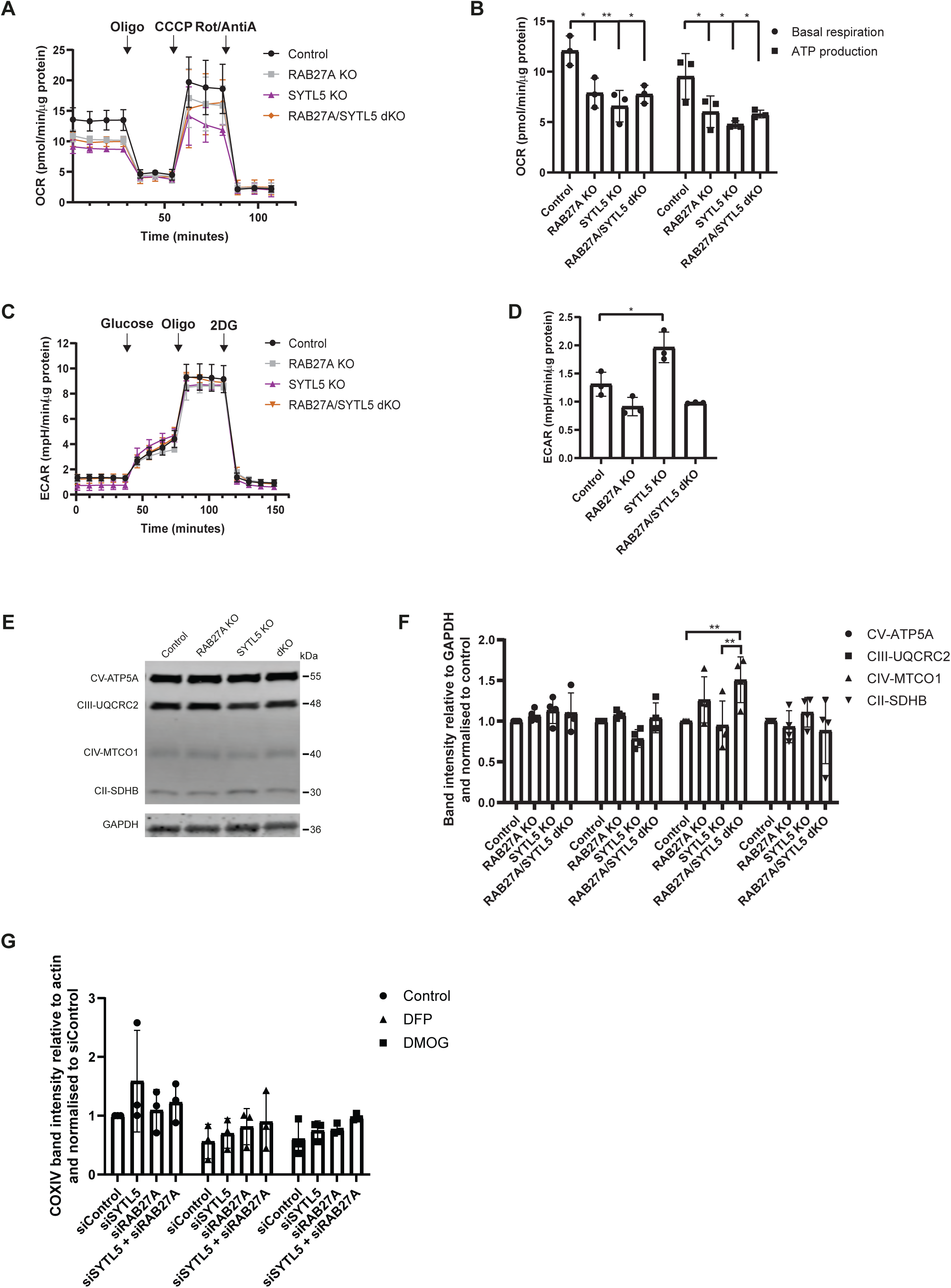
SYTL5 and RAB27A regulate mitochondrial bioenergetics. A. Mitochondrial oxygen consumption rate was analysed in U2OS control cells, SYTL5 KO, RAB27A KO and dKO cells using the Seahorse XF analyser. OCR was measured after sequential addition of oligomycin, CCCP and a combination of rotenone and antimycin-A into the assay medium. Error bars represent the mean with standard deviation between replicates (n=3). B. The four basal OCR measurements per well (before oligomycin addition) were averaged to obtain the basal OCR value and the non-mitochondrial respiration was subtracted to determine the basal respiration associated to each condition. The ATP production was calculated by subtracting the proton leak from the maximal respiratory capacity. Error bars represent the mean with standard deviation between replicates (n=3). Significance was determined by two-way ANOVA followed by Bonferroni multiple comparison test, * = p< 0.05, ** = p<0.01. C. The extracellular acidification rate (ECAR) was analysed in U2OS control cells, SYTL5 KO, RAB27A KO and dKO cells using the Seahorse XF analyser. ECAR was measured upon addition of glucose, oligomycin and 2-DG using control and SYTL5 depleted cells. Error bars represent the mean with standard deviation between replicates (n=3). D. The glycolysis rate of each condition was calculated by subtracting the ECAR value after 2-DG treatment to the ECAR after addition of glucose (Glycolysis=ECAR after addition of glucose−ECAR after 2-DG treatment). Error bars represent the mean with standard deviation between replicates (n=3). Significance was determined by one-way ANOVA followed by Tukey multiple comparison test, * = p< 0.05. E. Cell lysates from U2OS control, RAB27A KO, SYTL5 KO and dKO cells were analysed by western blot for the indicated electron transport chain complex-II, -III, -V and -V proteins. GAPDH was used as a loading control. F. Quantification of data in E. Band intensities were quantified relative to GAPDH and normalised to control. Error bars represent the mean with standard deviation between replicates (n=4). Significance was determined by two-way ANOVA followed by Bonferroni multiple comparison test, ** = p<0.01. G. U2OS cells were transfected with 20 nM control siRNA, SYTL5 siRNA #1 or RAB27A siRNA #1 for 48 h followed by 24 h incubation with 1 mM DFP or 1 mM DMOG. Cell lysates were analysed by western blot for COXIV. Actin was used as a loading control. Error bars represent the mean with standard deviation between replicates (n=3).

To test whether the decreased respiration in cells depleted of SYTL5 and/or RAB27A is coupled to a shift to glycolysis, we performed Seahorse analysis of the extracellular acidification rate (ECAR) in cells upon the sequential addition of glucose (increases glycolytic capacity), oligomycin (inhibits mitochondrial ATP production), and 2-DG (a glycolysis inhibitor which causes an abrupt decrease in ECAR). Intriguingly, glycolysis was significantly increased in SYTL5 KO cells as compared to control **(Figure 5C-D)**, indicating a possible metabolic shift to compensate for the decrease in OCR observed when SYTL5 was depleted **(Figure 5A-B).** In contrast, ECAR was reduced in cells lacking RAB27A compared to the control, which may indicate an impairment in glucose uptake **(Figure 5C)** and therefore in glycolysis **(Figure 5D)**. After adding oligomycin the ECAR increased in all cell lines, but in RAB27A KO and dKO cell lines, the increase was reduced when compared to the control cell line **(Figure 5C-D).**

The clear reduction in mitochondrial respiration observed in cells lacking RAB27A and SYTL5 led us to investigate the abundance of various ETC proteins. Depletion of SYTL5 and/or RAB27A did not affect the overall abundance of ATP5A (complex V), UQCRC2 (complex III) or SDHB (complex II), but we observed a significant increase in the level of MTCO1 (complex IV) in cells lacking RAB27A **(Figure 5E-F).** There was however no significant difference in the level of COXIV, another subunit of the ETC complex IV, upon siRNA-mediated knockdown of SYTL5, RAB27A or both compared to siControl in basal condition or upon induction of mitophagy with DFP or DMOG **(Figure 5G)**.

Taken together, our data show that both RAB27A and SYTL5 are important for normal mitochondrial respiration and ATP production while SYTL5 seems to regulate a metabolic switch to glycolysis.

### Low SYTL5 expression is related to reduced survival for adrenocortical carcinoma patients

Since our data indicate that SYTL5 may be a regulator of the metabolic switch from oxidative phosphorylation to glycolysis (the Warburg effect) **(Figure 5A-D)** that is frequently associated with cancer cells, we decided to analyse publicly available expression datasets to explore a possible link between SYTL5 and cancer. Using bulk RNA expression data from healthy/normal tissue samples in the GTEx project, we found that SYTL5 is preferentially expressed in adrenal gland and various regions of the brain (**Figure 6A).** Adrenocortical carcinoma (ACC) is a rare type of aggressive cancer with poor prognosis and limited treatment options that develops in the adrenal cortex^23^. A comparison of gene expression levels^31^ in normal adrenal gland samples and ACC samples showed that SYTL5 mRNA expression is significantly reduced in tumour samples. **(Figure 6B).** Furthermore, patient survival analysis using data obtained from the TCGA-ACC cohort, indicates that low SYTL5 mRNA expression is associated with a significantly reduced survival of ACC patients **(Figure 6C)**, suggesting a possible tumour suppressor function for SYTL5 in ACC cancer progression. RAB27A expression levels were also reduced in tumor tissue compared to normal samples, although to a lesser extent than SYTL5 **(Figure 6D),** and overall patient survival in ACC patients did not show any relationship with RAB27A expression levels **(Figure 6E)**.

**Figure 6.**
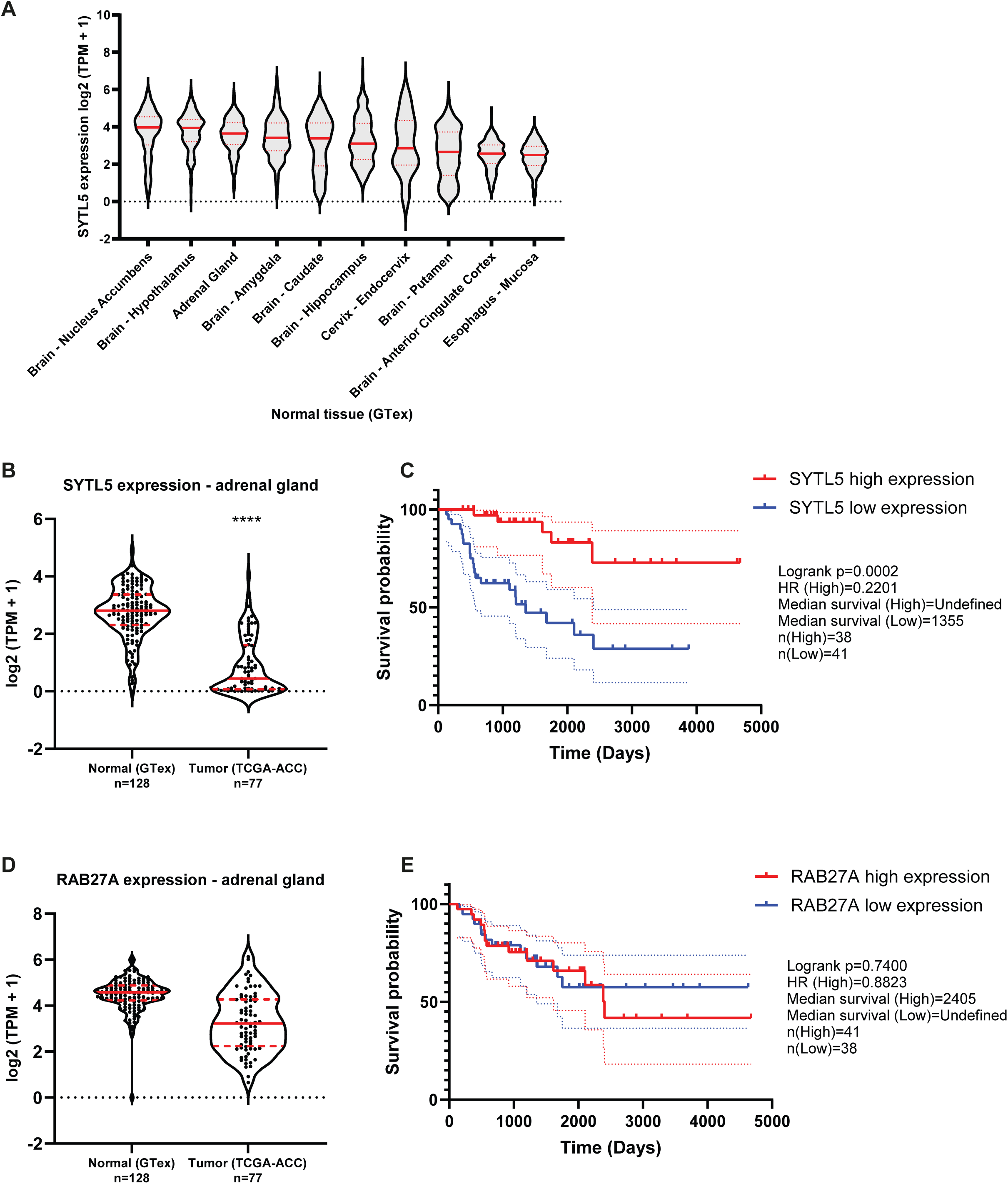
Low SYTL5 expression is related to reduced survival for adrenocortical carcinoma patients. A. SYTL5 expression levels in normal/healthy tissues, using data from the GTEx project (v8). Shown are the 10 tissue sites for which SYTL5 is most highly expressed (i.e. based on median expression levels). TPM = transcripts per million. Median is represented by a red solid line and quartiles by red dashed lines. B. Comparison of SYTL5 gene expression levels in normal adrenal gland samples (GTEx, n = 128) and adrenocortical carcinoma samples (The Cancer Genome Atlas (TCGA) ACC cohort, n = 77). Median expression is represented by a red solid line and quartiles by red dashed lines. Statistical analysis was performed using the unpaired t-test with Welch’s correction. TPM= transcripts per million. C. Kaplan-Meier plot for ACC patient survival related to SYTL5 expression levels. The events related to high SYTL5 expression (expression level above median) are indicated in red and the dotted lines represent 95% confidence interval. Events related to low SYTL5 expression (expression level below median) are indicated in blue and the dotted lines represent 95% confidence interval. Statistical analysis was performed using Logrank (Mantel-Cox) test. D. Comparison of RAB27A gene expression levels in normal adrenal gland samples (GTEx, n = 128) and adrenocortical carcinoma samples (The Cancer Genome Atlas (TCGA) ACC cohort, n = 77). Median expression is represented by a red solid line and quartiles by red dashed lines. Statistical analysis was performed using the unpaired t-test with Welch’s correction. E. Kaplan-Meier plot for ACC patient survival related to RAB27A expression levels. The events related to high RAB27A expression (expression level above median) are indicated in red and the dotted lines represent 95% confidence interval. Events related to low RAB27A expression (expression level below median) are indicated in blue and the dotted lines represent 95% confidence interval. Statistical analysis was performed using Logrank (Mantel-Cox) test.

The adrenal gland cortex secretes several steroid hormones, such as mineralocorticoids (e.g. aldosterone), glucocorticoids (e.g. cortisol), and adrenal androgens (DHEA and DHEA-sulphate)^32^, to the bloodstream. ACC tumours are characterised by excess adrenocortical hormone production in nearly 45–70% of patients, with hypercortisolism being the most common^23^. As mitochondria contain enzymes essential for steroid hormone biosynthesis from cholesterol^24^, we asked whether SYTL5 may regulate cortisol production. SYTL5 or RAB27A were depleted by siRNA in the ACC cell line H295R, followed by quantification of cortisol secretion using a colorimetric assay. We did not observe a significant change in cortisol secretion upon SYTL5 or RAB27A knockdown (**Supplementary Figure 6**), indicating that the correlation between low SYTL5 expression levels and ACC patient survival likely is unrelated to effects on cortisol production.

## Discussion

SYTL5 has been identified as a RAB27A effector protein with affinity for membranes through its C2 lipid-binding domains^1^, but nothing is known about a potential role of SYTL5 in regulation of RAB27A-dependent membrane trafficking pathways. Using the osteosarcoma derived U2OS cell line, here we confirm that SYTL5 interacts with RAB27A, and show that it localises to mitochondria and small, highly mobile vesicles interacting with the mitochondrial network, as well as to endolysosomal and endocytic compartments, indicating a role of SYTL5 in cellular membrane trafficking events.

Most interestingly, mitochondrial recruitment of SYTL5 involves coincidence detection, as it requires both its RAB27-binding SHD domain and its lipid-binding C2 domains. Indeed, we demonstrate that SYTL5 is recruited to mitochondria in a RAB27A-dependent manner and that RAB27A itself is localised to mitochondria in a manner dependent on its GTPase activity, as both RAB27A GDP and GTP mutants (T23N and Q78L, respectively) showed a dispersed cytoplasmic localisation. This is in line with a study showing that the GTPase activity of RAB27A is required for its localisation to melanosomes^33^. The mechanisms involved in mitochondrial recruitment of RAB27A are not clear. It is however interesting to note that other RAB proteins are recruited to the mitochondria by regulation of their GTPase activity. During parkin-mediated mitophagy, RABGEF1 (a guanine nucleotide exchange factor) is recruited through its ubiquitin-binding domain and directs mitochondrial localisation of RAB5, which subsequently leads to recruitment of RAB7 by the MON1/CCZ1 GEF complex^34^. Intriguingly, ubiquitination of the RAB27A GTPase activating protein alpha (RAB27B) is reduced in the brain of Parkin KO mouse compared to controls^35^, suggesting a possible connection of RAB27A with regulatory mechanisms that are linked with mitochondrial damage and dysfunction.

SYTL5 and RAB27A are both members of protein families, suggesting possible functional redundancies from Rab27B or one of the other SYTL isoforms. While RAB27B has a very low expression in U2OS cells, all five SYTL’s are expressed. However, when knocking out or knocking down SYTL5 and RAB27A we observe significant effects that we presume would be negated if their isoforms were providing functional redundancies. Moreover, we did not detect any other SYTL protein or RAB27B in the SYTL5 interactome, confirming that they do not form a complex with SYTL5.

Given the presence of mitochondria-associated small and highly mobile SYTL5/RAB27A-positive vesicles that contain mitochondrial material and stain positive for LAMP1, we speculate that RAB27A and SYTL5 facilitate mitophagy-mediated lysosomal delivery of mitochondria or selected mitochondrial components. Indeed, SYTL5 and RAB27A positive vesicles containing mitochondrial material were observed in cells upon hypoxia or hypoxia-mimicking (DFP and DMOG) treatments, where some vesicles stained positive for the autophagy markers p62 and LC3B upon co-treatment with the lysosomal V-ATPase inhibitor BafA1. Intriguingly, in cells treated with DFP and DMOG we observed a significant decrease in lysosomal cleavage of the mitophagy reporter pSu9-Halo-mGFP^36^ upon depletion of SYTL5 and/or RAB27A. Additionally, when blotting for various endogenous proteins of the electron transport chain complexes, we found that the level of the cytochrome c oxidase (complex IV) subunit MTCO1/COX1 was significantly increased in cells lacking RAB27A. However, neither the basal level of COXIV, another complex IV subunit, nor the lysosomal targeting of a mitochondrial matrix reporter (IMLS) were affected by depletion of SYTL5 and/or RAB27A. Expression of the mitophagy receptor BNIP3L, a HIF-1a target, was not affected by SYTL5 and/or RAB27A depletion. Mitochondrial recruitment of SYTL5 and RAB27A was also independent of the core autophagy machinery components ULK1 and VPS34. Taken together, our data indicate that mitochondrial RAB27A/SYTL5 function as positive regulators of selective clearance of mitochondrial components, in a piece-meal mitophagy-dependent manner.

The role of SYTL5 and RAB27A in the turnover of proteins involved in mitochondrial oxidative phosphorylation and ATP production is reminiscent of the recently described VDIM (Vesicles Derived from the Inner Mitochondrial membrane) pathway^16^. VDIMs are formed by IMM herniation through pores in the outer mitochondrial membrane, followed by their engulfment by lysosomes in proximity to mitochondria in a process that is independent of the core autophagy machinery. However, in contrast to our findings, VDIMs lack LC3 and p62 and oxidative stress-induced VDIM formation leads to selective degradation of IMM proteins, including COX4.

Mitochondria are known stress sensors and hubs for cellular adaptation^8^. When analysing the interactome of SYTL5, we found that many candidates were related to the cellular response to stress and oxygen-containing compounds, as well as vesicle-mediated transport and secretion. In line with a role for SYTL5/RAB27A in the regulation of mitochondrial stress, we found that cells lacking SYTL5 and/or RAB27A displayed reduced OXPHOS activity and ATP-production. A shift from OXPHOS to anaerobic glycolysis occurs in normal cells during hypoxic conditions and often in cancer cells even under normoxic conditions, commonly referred to as the Warburg effect^22^. Intriguingly, while the depletion of SYTL5 triggered a shift to glycolysis as observed by an increase in the extracellular acidification rate (ECAR) due to cellular lactate production, depletion of RAB27A or both RAB27A/SYTL5 had no significant effect, indicating that SYTL5 may function as a negative regulator of RAB27A and subsequently the Warburg effect.

The observed switch from OXPHOS to glycolysis in U2OS cells depleted of SYTL5 prompted us to look at cancer patient databases. We observed, using data from GTEx, that the adrenal gland is among the tissues with highest levels of SYTL5 gene expression, and that its expression is reduced in adrenocortical carcinoma samples (TCGA-ACC) when compared to normal adrenal gland tissue. Moreover, low SYTL5 expression is associated with reduced ACC patient survival. One characteristic of this type of cancer is an increase in steroid hormone production and secretion. Most steroid hormones are produced in mitochondria^24^ but we did not observe any changes in the level of cortisol in ACC cells depleted of SYTL5. The mechanisms linking SYTL5 to the Warburg effect and ACC need further investigation, but it is interesting to note that RAB27A expression levels do not correlate with ACC survival.

To summarise, we show that SYTL5 is a RAB27A effector protein being recruited to mitochondria in a RAB27A-dependent manner. Upon hypoxia, RAB27A and SYTL5 promote lysosomal clearance of selected mitochondrial components, and both proteins are needed for mitochondrial respiration. We identify SYTL5 as a negative regulator of the Warburg effect and potentially tumorigenesis.

## Materials and methods

### Antibodies and dyes

| Antibody | Source | Identifier |
| --- | --- | --- |
| α-Tubulin | Sigma-Aldrich | #T5168 |
| β-Actin | Cell signalling technology | #3700 |
| BNIP3L | Cell signalling technology | #12396 |
| COXIV | Cell signalling technology | #4850 |
| EGFP | Abcam; Clontech | #ab290; #632569 |
| FLAG Tag | Cell signalling technology | #14793 |
| GAPDH | Cell signalling technology | #5174 |
| Halo Tag | Promega | #G9211 |
| LAMP1 | Santa Cruz Biotechnology | #sc-20011 |
| LC3 | MBL international | #PM036 |
| OXPHOS | Abcam | #ab110413 |
| p62 | Progen | #GP62-C |
| RAB27A | Santa Cruz Biotechnology | #sc-74586 |
| SYTL5 | Sigma-Aldrich | #HPA026074 |
| TIM23 | BD biosciences | #611223 |
| TOMM20 | Santa Cruz Biotechnology | #sc-17764 |
| LysoTracker red DND-99 | ThermoFisher Scientific | #L7528 |
| MitoTracker™ Deep Red FM | ThermoFisher Scientific | # M22426 |
| MitoTracker™ Green FM | ThermoFisher Scientific | #M7514 |
| MitoTracker™ Red CMXRos | ThermoFisher Scientific | #M7512 |

#### Cell culture, lentivirus production and stable cell line generation

Human bone osteosarcoma epithelial cells (U2OS FlpIn TRex cell line, kindly provided by Steve Blacklow, Harvard Medical School, US) were grown in complete medium of Dulbeccós Modified Eagle Medium (DMEM; Lonza #12-604F and Gibco # 31966047) with 10% v/v foetal bovine serum (FBS; Sigma-Aldrich #F7524),100 U/ml Penicillin and 100 μg/ml Streptomycin (P/S; Thermo Fisher Scientific #15140122) at 37 °C with 5% CO2.

For lentiviral particle production HeK-FT cells were co-transfected with 1.6 μg of pLVX lentiviral vector, pCMV-VSV-G and psPAX2 using X-tremeGENE™ 9 DNA Transfection Reagent (Sigma #XTG9-RO). For virus infection, cells were seeded in complete medium and supplemented with 8 μg/ml polybrene (Santa Cruz Biotechnology #sc-134220). 2 μg/ml Puromycin (Sigma Aldrich #P7255) was added after 24 h post-infection for selection. To generate U2OS stably expressing SYTL5-EGFP, U2OS FlpIn TRex cells and U2OS FlpIn TRex_mScarlet-RAB27A were co-transfected with 0.1 μg pcDNA™5/FRT/TO_SYTL5-EGFP and 0.9 μg pOG44 Flp recombinase expression vector using Lipofectamine™ 2000 (Thermo Fisher Scientific #11668019) according to the manufacture’s instructions. To produce stable cell lines expressing pLVX-SV40-mScarlet-RAB27A(T23N/Q78L) and pLVX-CMV-SYTL5-EGFP-3xHA, U2OS dKO cells were infected with lentiviral particles followed by fluorescence-activated cell sorting (FACS) for double mScarlet/EGFP expressing cells. To generate stable cells expressing pLVX-CMV-SYTL5-EGFP-3xFLAG, pLVX-CMV-SYTL5 (ΔC2AB)-EGFP-3xFLAG and pLVX-CMV-SYTL5 (ΔSHD)-EGFP-3xFLAG, U2OS FlpIn TRex cells were infected with lentiviral particles. To generate U2OS stably expressing pSu9-Halo-mGFP, U2OS cells were infected with retroviral particles generated using pUMVC (addgene # 8449) and pMRX-IB-pSu9-HaloTag7-mGFP (addgene # 184905). Stable cell lines were maintained in complete medium with additional 5 μg/ml blasticidin S (Thermo Fisher Scientific #R21001) or 2 μg/ml puromycin (Sigma Aldrich #P7255). For starvation experiments, cells were washed with PBS and incubated for 4 h in Earlés balanced salt solution (EBSS; Gibco #24010043).

NCI-H295R cells were obtained from ATCC (#CRL-2128) and maintained in DMEM: F12, HEPES medium (Gibco #11330032) supplemented with 2.5% Nu-Serum (Corning #355100), 1% ITS+Premix Universal culture supplement (Corning #354352) and 100 μg/ml P/S (Thermo Fisher Scientific #15140122) at 37 °C with 5% CO2.

### Hypoxia and drug treatments

After cells were seeded, they were transferred to a INVIVO_2_ 200 Hypoxic workstation (Ruskinn). Hypoxia treatment was carried out for 24 hours with oxygen concentration set to 1%.

Drugs were added into the cell media. Treatment durations and drug concentrations are indicated in the figure legends. Drugs used were DFP (Sigma-Aldrich #379409), DMOG (Sigma-Aldrich #D3695), VPS34IN1 (Selleckchem # S7980) and MRT68291 (Selleckchem #S7949).

### Molecular cloning

SYTL5 was amplified from WT U2OS cDNA (5’-ATGTCTAAGAACTCAGAGTTCATC-3’ and 5’-TTAGAGCCTACATTTTCCCATG-3’) and cloned into Zero Blunt TOPO vector using Zero Blunt™ TOPO™ PCR Cloning Kit (Thermo Fisher Scientific #450245) according to the manufacture’s instructions. pcDNA™5/FRT/TO_SYTL5-EGFP was generated by Gibson Assembly using the TOPO-SYTL5 vector as a template for SYTL5 and cloned into a pcDNA™5/FRT Mammalian Expression Vector (Thermo Fisher Scientific #V601020) or into a pLVX-CMV lentiviral expression vector. Truncated SYTL5 (ΔSHD) and SYTL5 (ΔC2AB) were generated by Gibson assembly using the pcDNA™5/FRT/TO_SYTL5-EGFP construct as a template and cloned into a pLVX-CMV lentiviral expression vector.

RAB27A was amplified from WT U2OS cDNA and cloned into a pLVX-SV40 lentiviral expression vector. RAB27A mutations were performed using the site-directed mutagenesis QuikChange II kit (Agilent #200524) according to the manufacture’s instructions using the following primer pairs:

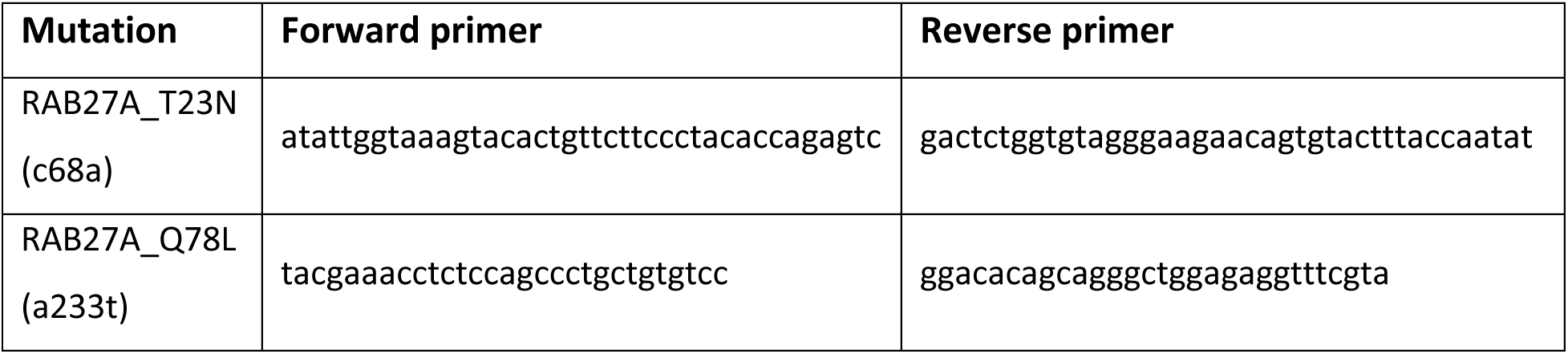

### CRISPR knockout cell line generation and validation

SYTL5 and RAB27A gene knockouts were performed as previously described by Ran *et al*., 2013. Briefly, gRNA target sequences were designed using Wellcome Sanger Institute Genome Editing (WGE) design tool available at https://wge.stemcell.sanger.ac.uk/find_crisprs.

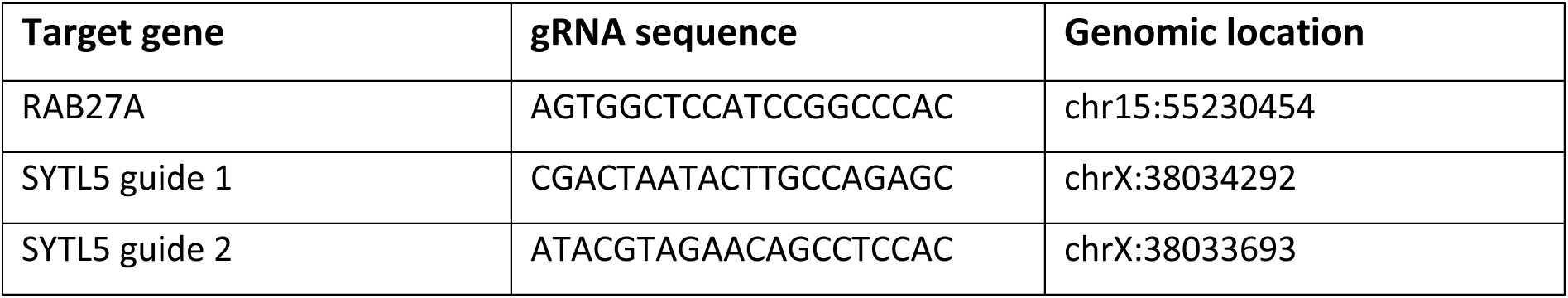

DNA oligos were annealed and phosphorylated before cloning into pSpCas9(BB)-2A-Puro (PX459) V2.0 (addgene # 62988). The obtained CRISPR plasmids were transfected into U2OS Flp-In TRex cells using Lipofectamine™ 2000 according to the manufacture’s instructions and selection with 2 μg/ml puromycin (Sigma Aldrich #P7255) was applied for 24 h. Limiting dilutions were performed using a concentration of 15 cells/ml and seeded in complete medium in a 384 well plate. Single colonies were collected ∼10 days after seeding and expanded for a further 3 weeks. For genotyping, gDNA from each single clone line was isolated using QIAamp DNA Mini Kit (QIAGEN #51304) and the targeted genomic region was amplified by PCR using flanking primers (RAB27A: 5’-ATAACCTCTCCCTTGACCTTGTATG-3’ and 5’-TAGATGCCTTTGGGATTTGTACTGA-3’; SYTL5: 5’-CAGGTCCCTTTCTTCTCGCA-3’ and 5’-TCAGTCAGCTGCAAGAGTGG-3’). The PCR product was cloned into Zero Blunt TOPO vector using Zero Blunt™ TOPO™ PCR Cloning Kit according to the manufacture’s instructions. RAB27A clonal lines were validated by western blot, and SYTL5 deletions were detected by running PCR products in 1% w/v agarose gel (Thermo Fisher Scientific #R2801).

### siRNA knockdown by reverse transfection

For siRNA knockdown, cells were incubated with OptiMEM (Thermo Fisher Scientific # 31985047) and Lipofectamine RNAiMAX (Thermo Fisher Scientific #13778150) for 5 min at room temperature (RT). Followed by addition of 20 nM siRNA diluted in OptiMEM and 15 min incubation at RT. Cells were fixed/harvested after 72 h of knockdown. The siRNA used for mRNA depletion are the following: siControl: 5’-UAACGACGCGACGACGUAAtt-3’; siSYTL5 #1: 5’-CUCUUAGAAGCAAAACGUAtt-3’; siSYTL5 #2: 5’-CAACAAGCGUAAGACCAAAtt-3’; siSYTL5 #3: 5’-GGUUUGUGCUUCAACCCAAtt-3’; siRAB27A #1: 5’-GGAAGACCAGUGUACUUUAtt-3’ and siRAB27A #2: 5’-CCAGUGUACUUUACCAAUAtt-3’.

### RNA isolation, cDNA synthesis and RT-PCR

RNA was isolated using TRIzol reagent (Thermo Fisher Scientific #15596026) and cDNA was synthesised using SuperScript III Reverse Transcriptase (Thermo Fisher Scientific #18080085) according to the manufacture’s instructions. qPCR analysis was performed using KAPA SYBR FAST qPCR Kit (Sigma-Aldrich #KK4601) and relative mRNA expression levels were normalised to the expression levels of TATA-binding protein (TBP) using the comparative ΔCt method. The primers used were the following:

SYTL5: 5’-GTGACAAAATCGCGCAGCTA-3’, 5’-GGACAACATCAGTGCCGAGA-3’

RAB27A: 5’-GGAGAGGTTTCGTAGCTTAACG-3’, 5’-CCACACAGCACTATATCTGGGT-3’

TBP: 5’-CAGAAAGTTCATCCTCTGGGCT-3’, 5’-TATATTCGGCGTTTCGGGCA-3’

### Immunofluorescent staining

Cells were fixed in 4% paraformaldehyde (PFA) (Sigma-Aldrich #158127) in PBS for at least 15min at 37°C, quenched for 10 min using 0.05 M NH4Cl in PBS, permeabilised in 0.05% saponin in PBS for 5 min and blocked in 1% BSA in PBS for 30 min at room temperature. Antibody staining was performed in a wet chamber for 1 h at room temperature and the antibodies were diluted in PBS containing 0.05% saponin. Nuclei staining was performed using Hoechst 33342 (Thermo Fisher Scientific #H1399) diluted in PBS at 1µg/ml.

### Light microscopy

Cells expressing inducible fluorescent proteins were induced with 100 ng/ml doxycycline (Clontech #631311) for 24 h. For live cell confocal microscopy cells were incubated with indicated dyes prior to imaging. Imaging was carried out with either the Andor Dragonfly High Speed Confocal Microscope or the Nikon Crest V3 with a 60x oil immersion objective (NA 1.4). High content widefield imaging was conducted with the ImageXpress Micro Confocal (Molecular Devices) microscope with a 20x objective (NA 0.45). Fixed immunofluorescence confocal imaging was performed using the Andor Dragonfly High Speed Confocal Microscope or Nikon Crest V3 both with a 60x oil immersion objective (NA 1.4) or Zeiss LSM 800 microscope (Zen Black 524 2012 SP5 FP3, Zeiss) with a 63x oil immersion objective (NA 1.4).

### Correlative light electron microscopy

U2OS cells co-expressing SYTL5-EGFP and mScarlet-RAB27A were seeded in gridded glass-bottom cell culture dishes (MatTek #P35G-1.5-14-CGRD). Cells growing in monolayer were fixed in warm (≈37°C) 3.7% PFA in 0.2 M HEPES (pH 7) and imaged using the Andor Dragonfly High Speed Confocal Microscope. After imaging, the samples were fixed using 2% v/v glutaraldehyde in 0.2 M HEPES, pH 7.4. After post-fixation in 2% v/v osmium tetraoxide and 3% v/v K4[Fe(CN)6], samples were embedded in Epon and 60 nm thickness sections were cut with a diamond knife. Sections were analysed using a JEM-1400 Plus Transmission Electron Microscope.

### Western blot

Cells were harvested in RIPA buffer (50 mM Tris pH 7.4, 150 mM NaCl (Merck #1064041000), 0,5% sodium deoxycholate (Sigma-Aldrich #30970), 1% NP-40 (Sigma-Aldrich #I3091), 1 mM EDTA (VWR #20302-293), 0,1% SDS (Sigma-Aldrich #L3771) containing 1x protease inhibitor (Merck #11697498001) and 1x phosphatase inhibitor (Merck #4906845001). 20 µg of lysate was loaded on SDS-PAGE gel (BioRad #5671095) and proteins transferred to a PVDF membrane (Merck #IPFL00010) which was blocked using 5x v/v casein blocking buffer in PBS for 1 h at room temperature. The obtained membranes were immunoblotted using primary and secondary antibodies with washing steps using PBST (1x Phosphate-Buffered Saline with 1% Tween20 (Sigma-Aldrich #P1379)). Membranes were analysed using the Odyssey CLx imaging system (Li-cor biosciences) and protein levels were quantified by densitometry with ImageStudio Lite software (Li-cor biosciences).

### Macro-mitophagy analysis

siRNA knockdown was conducted as previously described in U2OS-IMLS cells seeded in complete media supplemented with 100 ng/ml doxycycline. Culture media was replaced 24 h after transfection and 1 mM DFP (Sigma-Aldrich #379409) treatment was added after 48 h. For U2OS-IMLS cells expressing PARKIN, 20 μM CCCP (Enzo Life Sciences #BML-CM124-0500) was used for 16 h and 10 μM Q-VD-OPh (Sigma-Aldrich #SML0063-1MG) pan-caspase inhibitor was included to support cell survival^11,37^. Control wells with 100 nM BafA1 (AH diagnostics #BML-CM110-0100) were dosed 2 h prior to fixation. After 72 h, cells were washed with PBS pH 7 and fixed in warm (≈37°C) 3.7% PFA in 0.2 M HEPES (pH 7) for 15 min at 37°C. After washing with PBS, cells were incubated with 2 μg/ml Hoechst and widefield images were obtained using a high-content imaging microscope. The area of red-only puncta per cell (which represent mitochondria delivered to lysosomes as the EGFP signal is quenched in the acidic lysosome) was quantified using a Cell Profiler pipeline. Addition of BafA1 was used as a control to confirm that the mCherry-only puncta corresponded to lysosomal structures.

### Halo assay for mitophagy flux

siRNA knockdown was conducted as previously described in U2OS cells expressing the pSu9-Halo-mGFP reporter. 48 h after transfection culture media was changed to contain 100 nM tetramethylrhodamine (TMR)-conjugated Halo ligand (Promega #G8251). After 20 min incubation, cells were washed twice with PBS and the media replaced with either media containing no treatment, 1 mM DFP or 1 mM DMOG. 72 h after transfection cells were harvested and western blot conducted (as described earlier). Membranes were incubated with the Halo Tag antibody and actin antibody. Membranes were imaged using the Chemidoc MP system (Bio-Rad) and band intensity quantified in Image Lab 6.1 (Bio-Rad). Bands were normalised to actin loading control. The percentage cleaved Halo was calculated by the formula (Free Halo / (Free Halo + Full Length)) x 100.

### Oxygen consumption rate measurement by Seahorse XF analyser

U2OS cells were seeded in complete medium at the density of 4x10^4^ cells/well into XF24 cell culture microplates (Agilent #100777-004). The sensor cartridge was hydrated for 24 h using XF calibrant in a 37°C non-CO_2_ incubator. Before analysis, cell medium was replaced by warm (≈37°C) DMEM without Sodium Bicarbonate (pH 7.4) and placed for 1 h in a 37°C non-CO_2_ incubator. Each measurement cycle consisted of a mixing time of 3 min and data acquisition period of 3 min (three data points). OCR results refer to the average rates during each measurement cycle. The final concentration for each mitochondrial inhibitor used in this assay was 1.5 µM Oligomycin A (SelleckChem #S1478), 1 µM CCCP, 0.5 µM Rotenone (Sigma Aldrich #R8875-1G) and 0.5 µM Antimycin (Sigma Aldrich #A8674). Four baseline measurements were performed before adding each compound and three response measurements were taken after each compound was added. Data analysis was performed using Seahorse Analytics software available at https://seahorseanalytics.agilent.com/ and OCR data was normalised to protein concentration in each well.

### Extracellular acidification rate (ECAR) measurement by Seahorse XF analyser

U2OS cells were seeded in complete medium at the density of 4x10^4^ cells/well into XF24 cell culture microplates. The sensor cartridge was hydrated for 24 h using XF calibrant in a 37°C non-CO_2_ incubator. Before analysis, cell medium was replaced by warm (≈37°C) XF DMEM media (Seahorse Bioscience #102353-100) supplemented with 2 mM L-Glutamine (BioNordika #BE17-605E) and pH adjusted to 7.4. The final concentration for each reagent used in this assay was 11 mM Glucose, 1.3 µM Oligomycin A, 0.1 M 2-Deoxy-Glucose (2-DG; Sigma Aldrich #D8375-1G)^38^. Each measurement cycle consisted of a mixing time of 3 min and data acquisition period of 3 min (four data points). ECAR results refers to the average rates during each measurement cycle. Five baseline measurements were performed before adding each compound and four response measurements were taken after each compound was added. Data was normalised to protein concentration using Seahorse Analytics software available at https://seahorseanalytics.agilent.com/ and the glycolysis rate was calculated by subtracting the ECAR value after 2-DG treatment from the ECAR value after addition of glucose.

### Purification of crude mitochondrial fraction

To purify crude mitochondrial fraction, cells expressing EGFP (control), mScarlet-RAB27A, SYTL5-EGFP and SYTL5(ΔC2AB)-EGFP were harvested from one 150 mm (80-90% confluency) dish, washed twice with ice cold homogenisation buffer (210 mM mannitol (Sigma Aldrich #M4125), 70 mM sucrose (Sigma Aldrich #S0389), 5 mM HEPES pH 7.12 at 25°C) and harvested in 1 mL homogenisation buffer supplemented (HBS) with 1 mM EGTA (Sigma Aldrich #E3889) and 1x protease inhibitor. The obtained cell lysate was further mechanically homogenised at 4°C using a cell homogeniser (Isobiotech) equipped with a 16 µm clearance ball by repeatedly passing the cell lysate for 10x repeats with the aid of 2 disposable syringes. The homogenised solution was further spun down at 1500xg for 3 min at 4°C to collect the TCL and the obtained supernatant collected and spun down at 13,000xg for 20 min. The obtained supernatant corresponds to the cytosolic fraction and the pellet was resuspended in 2 mL HBS and spun down at 13,000xg for 20 min. The obtained pellet corresponds to the enriched mitochondrial fraction and was resuspended in 900 µL HBS. 50 µL of each cellular fraction was resuspended in 30 µL 2x SDS-sample buffer and boiled at 100°C for 5 min and further analysed using SDS-PAGE.

### GFP-TRAP and FLAG immunoprecipitation

Cells expressing EGFP were harvested from a 100 mm dish (80-90% confluency), washed with ice cold PBS and scraped using ice cold isolation buffer (1% NP-40, 2 mM EDTA, 136 mM NaCl, 20 mM Tris-HCl pH 7.4, 10% glycerol, 1x protease inhibitor and 1x phosphatase inhibitor). Cell lysate was spun down at 20,000xg for 10 min and the supernatant added to 25 µL equilibrated GFP-trap agarose beads (Chromotek #gta-20). Samples were rotated end-over-end for 1 h at 4°C. After 4x washing steps, the beads were collected by centrifugation at 2700xg for 1 min, resuspended in 30 µL 2x SDS-sample buffer and boiled at 100 °C for 5 min. The obtained supernatant was analysed by SDS-PAGE.

U2OS cells expressing pLVX-CMV-SYTL5-EGFP-3xFLAG, pLVX-CMV-SYTL5(ΔC2AB)-EGFP-3xFLAG and pLVX-CMV-SYTL5(ΔSHD)-EGFP-3xFLAG were harvested from a 150 mm dish (80-90% confluency), washed 2x with cold PBS and scraped using 1 ml of cold lysis buffer (50 mM Tris-HCl, pH 7.4,150 mM NaCl, 1% Triton X-100, 5 mM CaCl_2_ (Merck #1023821000, and 1x protease inhibitor. Cell lysates were spun down at 20,000xg for 10 min and the resulting supernatants added to 40 µL of pre-washed anti-FLAG M2-Agarose Affinity Gel (Sigma Aldrich #FLAGIPT-1). Samples were rotated end-over-end for 2 h at 4°C. After 3x washing steps, the affinity gel was centrifuged at 5000xg for 30 seconds and the FLAG fusion proteins eluted using 3x FLAG peptide according to manufacturer’s protocol. The eluted FLAG fusion proteins were diluted in 4 ml 1x TBST supplemented with 3% BSA and 5 mM CaCl_2_.

### I*n-vitro* protein-lipid binding assay

Eluted FLAG fusion proteins (SYTL5-EGFP-3xFLAG, SYTL5(ΔC2AB)-EGFP-3xFLAG and SYTL5(ΔSHD)-EGFP-3xFLAG were diluted in 1x TBST supplemented with 3% BSA and 5 mM CaCl_2_. PIP strips (Echelon Biosciences #P-6001) and membrane lipid strips (Echelon Biosciences #P-6002) were blocked in 1x TBST supplemented with 3% BSA for 1 h at room temperature and incubated with the diluted FLAG fusion protein for 1 h at 4°C with gentle agitation. Lipid membranes were washed 3x with TBST and incubated with FLAG Tag primary antibody for 1 h at room temperature and detected using SuperSignal West Dura Extended Duration Substrate (Thermo Fisher Scientific #34076).

### LC-MS/MS analysis of SYTL5-EGFP interactome

Cells expressing SYTL5-EGFP or EGFP (control) were harvested using NP-40 lysis buffer (10 mM Tris-HCl pH 7.5, 150 mM NaCl, 0.5 mM EDTA, 0.5% NP-40, 1x protease inhibitor. SYTL5-EGFP was immunoprecipitated using GFP-Trap agarose beads according to the manufacturer’s protocol. The obtained beads were washed twice with 50 mM ammonium bicarbonate, reduced, alkylated and further digested by trypsin for overnight at 37 °C. Digested peptides were transferred to a new tube, acidified and the peptides were de-salted for MS analysis.

Peptide samples were dissolved in 10 µl 0.1% formic buffer and 3 µl was loaded for MS analysis. LC-MS/MS analysis of the resulting peptides was performed using an Easy nLC1000 liquid chromatography system (Thermo Electron) coupled to a QExactive HF Hybrid Quadrupole-Orbitrap mass spectrometer (Thermo Electron) with a nanoelectrospray ion source (EasySpray, Thermo Electron). The LC separation of peptides was performed using an EasySpray C18 analytical column (2 µm particle size, 100 Å, 75 μm inner diameter and 25 cm (Thermo Fisher Scientific). Peptides were separated over a 90min gradient from 2% to 30% (v/v) ACN in 0.1% (v/v) FA, after which the column was washed using 90% (v/v) ACN in 0.1% (v/v) FA for 20 min (flow rate 0.3 μL/min). All LC-MS/MS analyses were operated in data-dependent mode where the most intense peptides were automatically selected for fragmentation by high-energy collision-induced dissociation.

Raw files from LC-MS/MS analyses were submitted to MaxQuant 1.6.17.0 software^39^ for peptide/protein identification. Parameters were set as follow: Carbamidomethyl (C) was set as a fixed modification and PTY; protein N-acetylation and methionine oxidation as variable modifications. First search error window of 20 ppm and mains search error of 6 ppm. Trypsin without proline restriction enzyme option was used, with two allowed miscleavages. Minimal unique peptides were set to one, and FDR allowed was 0.01 (1%) for peptide and protein identification. The Uniprot human database was used. Generation of reversed sequences was selected to assign FDR rates. Further analysis was performed with Perseus^40^, limma/voom^41^ and Package R^42^. Volcano plots were plotted with EnhancedVolcano^43^. GO analysis was performed with shinyGO^44^.

### Bulk gene expression analysis – normal and tumor tissue

Bulk gene expression distribution in healthy/normal tissue samples was collected from GTEx v8 (PMID: 26484571). We utilised a combined RNA-seq expression dataset of TCGA and GTEx samples available through the Xena platform to investigate SYTL5 and RAB27A expression in normal and tumor tissue (PMID: 32444850). Here, data from a total of n = 77 samples provided normal adrenal gland expression (GTEx), and a total of n = 128 samples provided expression in adrenocortical carcinoma samples (TCGA-ACC). Survival data from samples in the TCGA-ACC cohort (one record per patient) was obtained using the TCGABiolinks R package^45^. Survival curve analysis was performed using GraphPad Prism 8.0.1 using Logrank test (Mantel-Cox).

### Cortisol assay

To measure cortisol in cell culture supernates, siRNA knockdown by reverse transfection with siSYTL5 and siRAB27A was conducted as described earlier in NCI-H295R cells. Cells were harvested with TNTE Lysis Buffer. Cortisol from lysates was measured using the R&D Systems® Cortisol Immunoassay kit (# KGE008B) as described in the protocol provided from the company. Controls were performed using cells treated with 50 uM Forskolin as a positive control to stimulate cortisol secretion^46^, or 10 uM mitotane to inhibit cortisol production^47^.

### Statistical analysis

Statistical analysis was performed with GraphPad Prism (8.0.1, 9.3.1 and 10.2.0 versions) and the tests used are indicated in the respective figure legends. All values come from distinct samples.

**** = p<0.0001, *** = p<0.001, ** = p<0.01, * = p< 0.05.

## Acknowledgments

We thank Jenni Lane for assisting with EM sample preparation, Sachin Singh and Tuula Nyman at the OUS Proteomics Core Facility for assisting with mass spectrometry-based proteomic experiments, Ankush Sharma for help with analysis of mass spectrometry data, Anna Lång at ALM Core Facility Gaustad for assisting with widefield microscopy experiments, Santosh Phuyal for experimental assistance and Serhiy Pankiv for providing the lentiviral mScarlet-RAB expression constructs. We also thank Patricia González-Rodríguez for providing us with the U2OS pSu9-HaloTag7-mGFP cells and Viola Nähse for her contributions and advice regarding CRISPR knock-in strategies.

This project was funded by the Research Council of Norway through its Centres of Excellence funding scheme (project number 262652) and FRIPRO grant (project number 249753), the Norwegian Cancer Society (project number 190251), Marie Skłodowska-Curie ETN grant under the European Union’s Horizon 2020 Research and Innovation Programme (Grant Agreement No 765912 DRIVE) and by the University of Oslo Scientia Fellow program through the MSC scheme – Co-funding of Regional, National and International Programmes (COFUND). Mass spectrometry-based proteomic analyses were performed by the Proteomics Core Facility, Department of Immunology, University of Oslo/Oslo University Hospital, which is supported by the Core Facilities program of the South-Eastern Norway Regional Health Authority. This core facility is also a member of the National Network of Advanced Proteomics Infrastructure (NAPI), which is funded by the Research Council of Norway INFRASTRUKTUR-program (project number: 295910).

## Author contributions

A.L., L.S.J., L.T.M. and A.S. designed and conceived the experimental work. A.L., L.S.J., L.T.M.,

M.Y.W.N. and S.J.R. performed the experiments and data analyses. S.S. carried out MS and GO term enrichment data analysis. S.N. provided GTEx and TCGA data analysis. E.L.E. performed and analysed CLEM data. A.L., L.S.J. and A.S. wrote the manuscript.

## Author declarations

The authors have nothing to declare.

---

**Supplementary video 1.** Live confocal microscopy imaging tracking SYTL5 vesicle movement along filaments positive for MitoTracker red.

**Figure S1.**
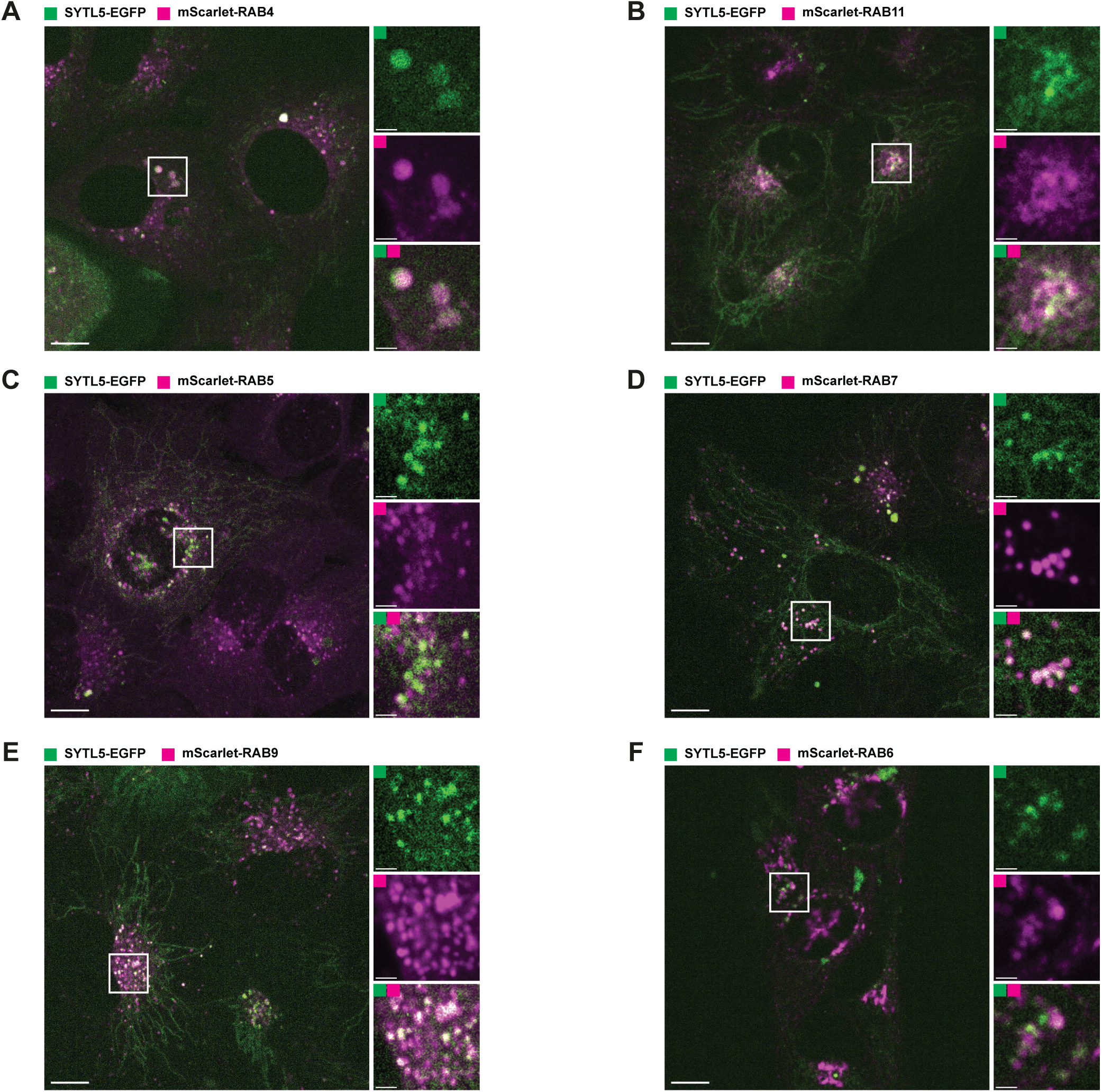
A-F. SYTL5-EGFP expression was induced for 24 h with 100 ng/ml doxycycline in U2OS cells with stable inducible expression of SYTL5-EGFP and constitutive expression of mScarlet-RAB4 (A), mScarlet-RAB11 (B), mScarlet-RAB5 (C), mScarlet-RAB7 (D), mScarlet-RAB9 (E) or mScarlet-RAB6 (F) followed by live confocal microscopy imaging analysis. Scale bars: 10 μm, 2 μm (insets).

**Figure S2.**
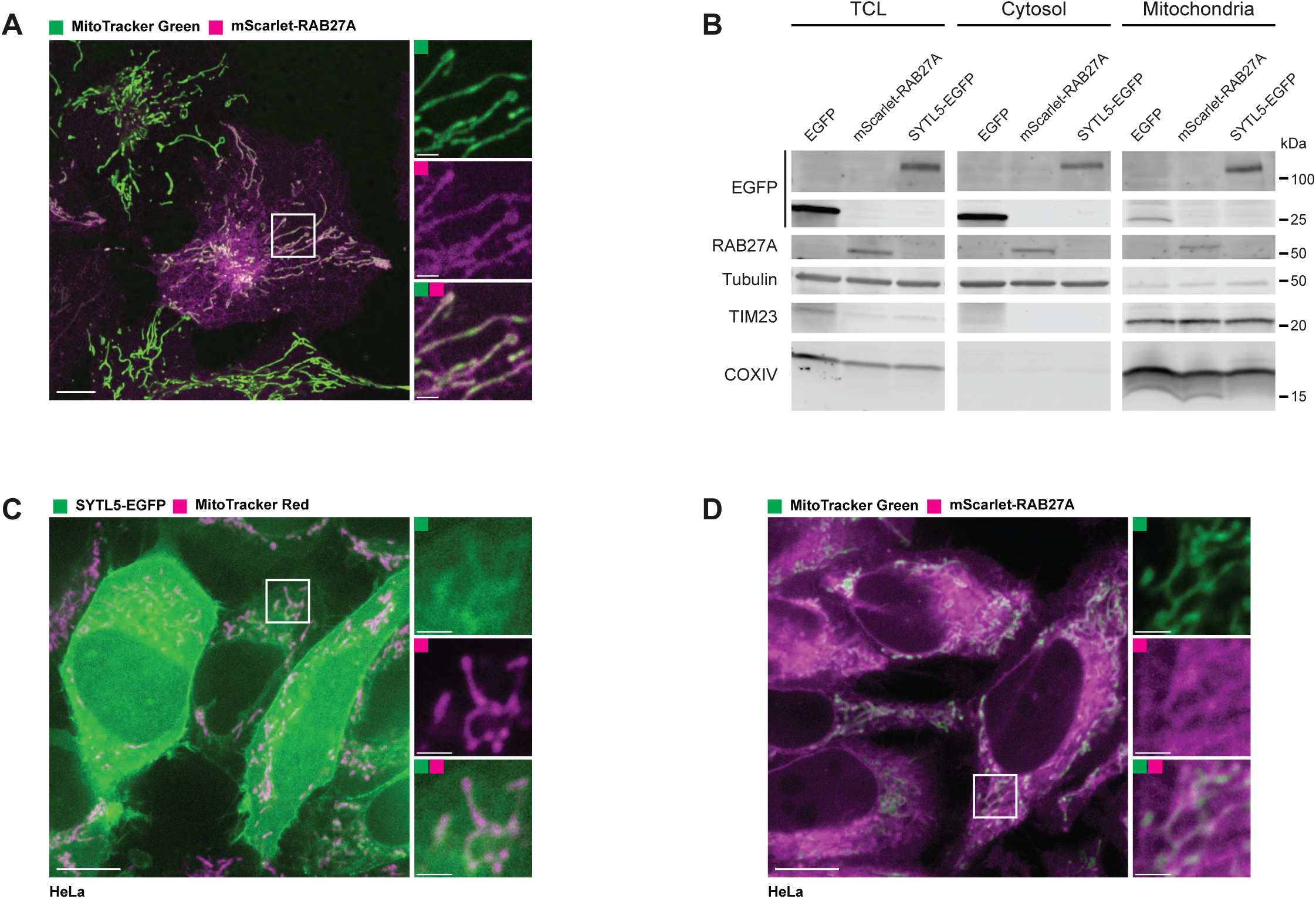
A. Live confocal microscopy imaging of U2OS constitutively expressing mScarlet-RAB27A co-stained with MitoTracker green added 30 min before imaging. Scale bars: 10 μm, 2 μm (insets). B. Total cell lysate (TCL), cytosol, and a crude mitochondria fraction from U2OS control cells and cells with stable expression of mScarlet-RAB27A or SYTL5-EGFP were analysed by western blotting for EGFP, RAB27A, tubulin, TIM23 and COXIV. C. Live confocal microscopy imaging of HeLa constitutively expressing SYTL5-EGFP co-stained with MitoTracker Red added 30 min before imaging. Scale bars: 10 μm, 2 μm (insets). D. Live confocal microscopy imaging of HeLa constitutively expressing mScarlet-RAB27A co-stained with MitoTracker green added 30 min before imaging. Scale bars: 10 μm, 2 μm (insets).

**Figure S3.**
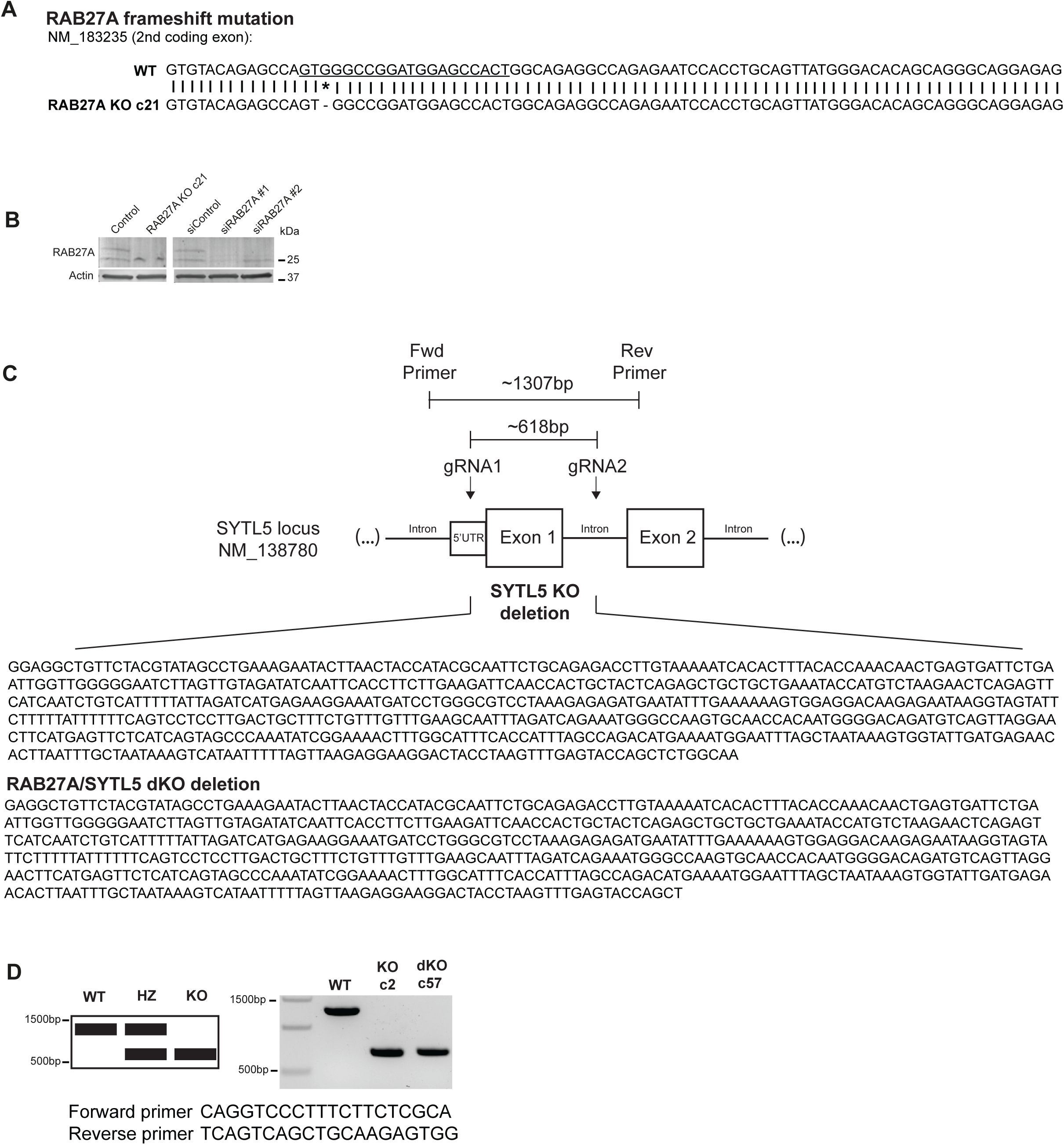
A. RAB27A KO validation by genotyping. The WT RAB27A gDNA sequence was aligned with the RAB27A KO clone 21(c21) gDNA sequence. CRISPR/Cas9 editing produced an indel (single nucleotide deletion indicated by *) in a genomic location shared between all RAB27A isoforms. gRNA targeting region is underlined in the WT RAB27A gDNA sequence. B. RAB27A KO validation by western blot. Cell lysates from U2OS control cells and RAB27A KO cells (c21) were compared to cell lysates where RAB27A was depleted for 72 h using 2 independent siRNAs and probed with RAB27 and actin antibodies. C. SYTL5 CRISPR/Cas9 exon deletion was performed using two gRNA simultaneously. These gRNA target the 5’UTR or intronic regions of SYTL5 (transcript NM_138780) that are shared by all SYTL5 isoforms. Genomic deletion validation and individual clone genotyping was performed using a primer pair that flanked the region to be edited (about 1307 bp length between these primer binding sites). D. SYTL5 KO validation was performed with conventional PCR reaction. A PCR product of 1307 bp is expected for wild-type U2OS gDNA, while a PCR product of about 618 bp is expected if a clone has a deletion in the targeted region. This method was used to validate the SYTL5 KO clone2 and for dKO RAB27A/SYTL5 clone57. The deleted sequence includes a coding exon from SYTL5, resulting in a non-functional SYTL5 protein.

**Figure S4.**
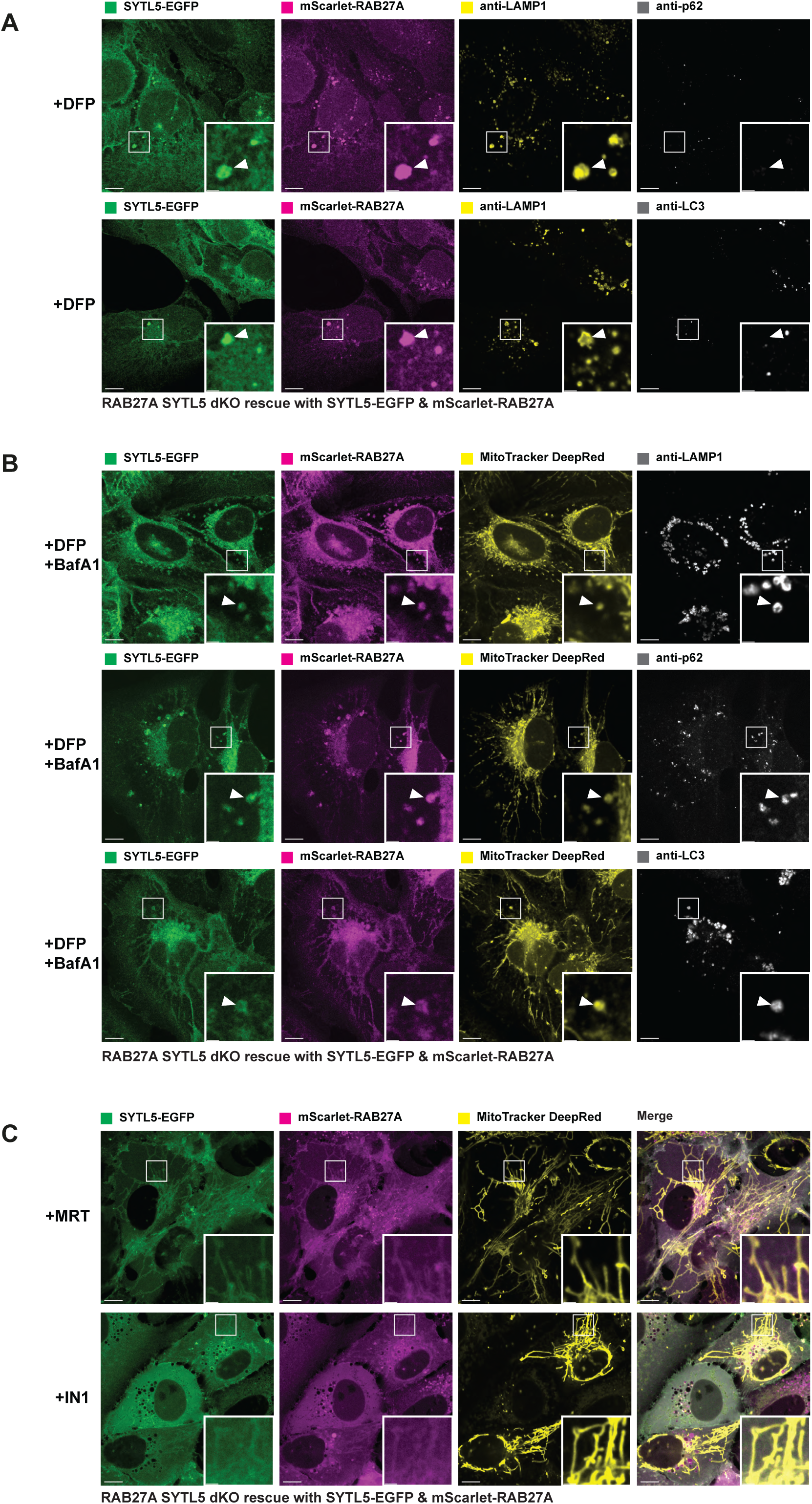
A. SYTL5/RAB27A dKO U2OS cells rescued with SYTL5-EGFP and mScarlet-RAB27A were treated with 1 mM DFP for 24 h. Fixed cells were stained with a LAMP1 antibody in addition to either p62 (upper panel) or LC3 (lower panels) antibodies and imaged by confocal microscopy. Arrowheads point to SYTL5/RAB27A-positive structures. Scale bars: 10 μm, 2 μm (insets). B. SYTL5/RAB27A dKO U2OS cells rescued with SYTL5-EGFP and mScarlet-RAB27A were treated with 1 mM DFP for 24 h and 100 nM BafA1 for the final 4 h. All cells were stained with 50 nM MitoTracker DR 30 min prior to fixing. Fixed cells were stained with either LAMP1 (upper panel), p62 (middle panel) or LC3 (lower panel) antibody and imaged by confocal microscopy. Arrowheads point to SYTL5/RAB27A-positive structures. Scale bars: 10 μm, 2 μm (insets). C. SYTL5/RAB27A dKO U2OS cells rescued with SYTL5-EGFP and mScarlet-RAB27A were treated with 1 μM MRT68921 for 2 h (upper panel) or 5 μM Vps34-IN1 for 2 h (lower panel). All cells were stained with 50 nM MitoTracker DR 30 min prior to live imaging confocal microscopy. Scale bars: 10 μm, 2 μm (insets).

**Figure S5.**
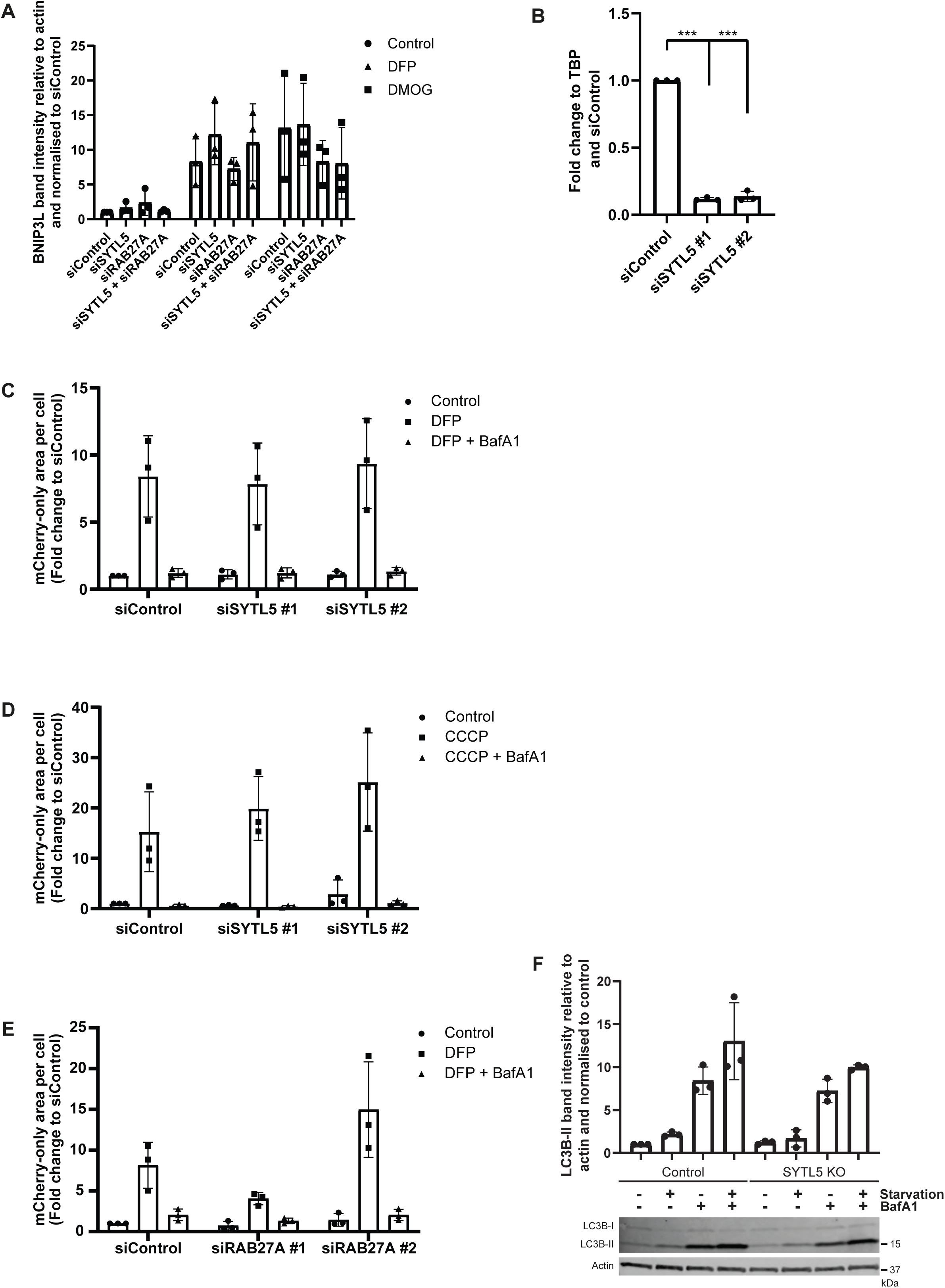
B. U2OS cells were transfected with 20 nM control siRNA, SYTL5 siRNA #1 or RAB27A siRNA #1 for 48 h followed by 24 h incubation with 1 mM DFP or 1 mM DMOG. Cell lysates were analysed by western blot for BNIP3L. Actin was used as a loading control. Error bars represent the mean with standard deviation between replicates (n=3). C. U2OS cells were transfected with 20 nM control siRNA or SYTL5 siRNA oligos for 72 h. Transfection efficiency was analysed by qPCR where the relative mRNA expression of SYTL5 was normalised to the expression levels of TBP and siControl. Error bars represent the standard deviation between replicates (n=3). Significance was determined by one-way ANOVA followed by Dunnett multiple comparison test, *** = p<0.001. D. U2OS cells expressing the mitochondrial matrix reporter NIPSNAP1^1–53^-EGFP-mCherry^29^ (referred to as IMLS cells) were transfected with 20 nM control siRNA or SYTL5 siRNA oligos for 48 h before 24 h incubation with 1 mM DFP in the absence or presence of 100 nM BafA1 for the last 2 h. After fixation, cell nuclei were stained with Hoechst and widefield images were obtained using a high-content imaging microscope. The area of red-only puncta per cell (>1000 cells analysed, represent mitochondria delivered to lysosomes as the EGFP signal is quenched in the acidic lysosome) was quantified using a Cell Profiler pipeline. The obtained results were normalised to the control siRNA and error bars represent the mean with standard deviation between replicates (n=3). Significance was determined by two-way ANOVA followed by Bonferroni multiple comparison test. E. U2OS IMLS cells expressing PARKIN were transfected with 20 nM control siRNA or SYTL5 siRNA oligos for 48 h before a 16 h incubation with 20 μM CCCP to induce mitophagy, with treatment with 100 nM BafA1 or not for the last 2 h. The area of red-only puncta per cell (>1000 cells analysed) was quantified using Cell Profiler from widefield images obtained using a high-content imaging microscope. The obtained results were normalised to the control siRNA and error bars represent the mean with standard deviation between replicates (n=3). Significance was determined by two-way ANOVA followed by Tukey’s multiple comparison test. F. U2OS IMLS cells were transfected with 20 nM control siRNA or RAB27A siRNA oligos for 48 h before 24 h incubation with 1 mM DFP in the absence or presence of 100 nM BafA1 for the last 2 h. After fixation, cell nuclei were stained with Hoechst and widefield images were obtained using a high-content imaging microscope. The area of red-only puncta per cell (>1000 cells analysed) was quantified using a Cell Profiler pipeline. The obtained results were normalised to the siControl, and error bars represent the mean with standard deviation between replicates (n=3). Significance was determined by two-way ANOVA followed by Bonferroni multiple comparison test. G. WT (control) or SYTL5 KO U2OS cells were starved in EBSS for 4 h or incubated with complete medium in the presence or absence of 100 nM BafA1 the last 2 h. Cell lysates were harvested and analysed by western blot for LC3B and actin proteins. LC3B-II band intensity was quantified relative to actin and normalised to control. Error bars represent the mean with standard deviation between replicates (n=3). Significance was determined by one-way ANOVA followed by Bonferroni multiple comparison test.

**Figure S6.**
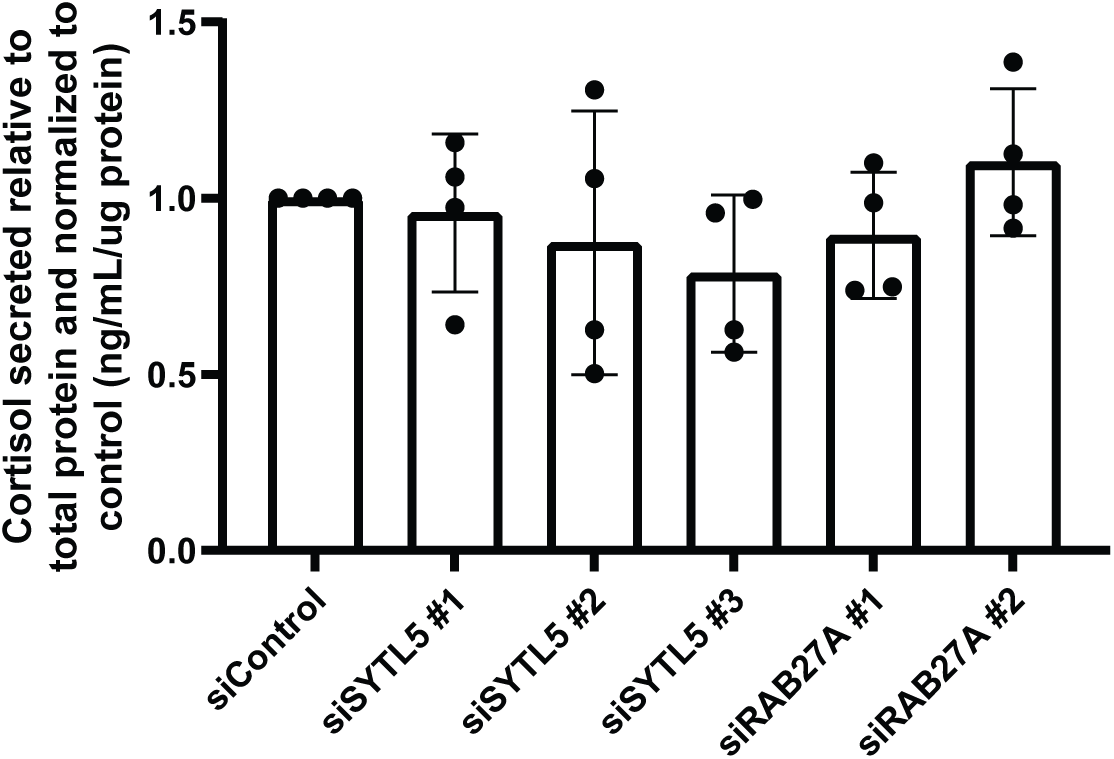
: NCl-H295R cells were transfected with 20 nM control siRNA, SYTL5 siRNA or RAB27A siRNA oligos for 72 h. Cortisol secretion was measured and normalised to siControl. Error bars represent the mean with standard deviation between replicates (n=4).

## Notes

### Competing Interest Statement

The authors have declared no competing interest.

### Summary of Updates

This version of our manuscript entitled The RAB27A effector SYTL5 regulates mitophagy and mitochondrial metabolism has been revised to update the following: New data has been included showing that the hypoxia-induced SYTL5-RAB27A positive vesicles containing mitochondrial material stain positive for the autophagy proteins p62 and LC3B (Supplementary Figure 4B) and that SYTL5 and RAB27A promote mitophagy of selected mitochondrial proteins in response to hypoxia (Figure 5G, and Supplementary Figure 5A). The discussion has been slightly revised in response to the reviewers' comments. Taken together, the data presented in this manuscript demonstrate that Synaptotagmin-Like 5 (SYTL5) is a RAB27A effector protein that localizes to mitochondria in a RAB27A-dependent manner and that both SYTL5 and RAB27A promotes mitophagy of selected mitochondrial proteins, as well as mitochondrial respiration. Intriguingly, cells lacking SYTL5 show increased glucose uptake and SYTL5 expression is significantly reduced in tumors of the adrenal gland and correlates with poor survival of patients with adrenocortical carcinoma.

## References

1 Kuroda, T. S., Fukuda, M., Ariga, H. & Mikoshiba, K. Synaptotagmin-like protein 5: a novel Rab27A effector with C-terminal tandem C2 domains. Biochem Biophys Res Commun 293, 899–906 (2002). 10.1016/s0006-291x(02)00320-0

2 Kuroda, T. S., Fukuda, M., Ariga, H. & Mikoshiba, K. The Slp homology domain of synaptotagmin-like proteins 1-4 and Slac2 functions as a novel Rab27A binding domain. J. Biol. Chem. 277, 9212–9218 (2002). 10.1074/jbc.M112414200

3 Nalefski, E. A. & Falke, J. J. The C2 domain calcium-binding motif: structural and functional diversity. Protein Sci. 5, 2375–2390 (1996). 10.1002/pro.5560051201

4 Gálvez-Santisteban, M. et al. Synaptotagmin-like proteins control the formation of a single apical membrane domain in epithelial cells. Nat. Cell Biol. 14, 838–849 (2012). 10.1038/ncb2541

5 Fukuda, M. The C2A domain of synaptotagmin-like protein 3 (Slp3) is an atypical calcium-dependent phospholipid-binding machine: comparison with the C2A domain of synaptotagmin I. Biochem. J. 366, 681–687 (2002). 10.1042/bj20020484

6 Yu, M. et al. Exophilin4/Slp2-a targets glucagon granules to the plasma membrane through unique Ca2+-inhibitory phospholipid-binding activity of the C2A domain. Molecular biology of the cell 18, 688–696 (2007). 10.1091/mbc.e06-10-0914

7 Fukuda, M. Rab27 effectors, pleiotropic regulators in secretory pathways. Traffic 14, 949–963 (2013). 10.1111/tra.12083

8 Montava-Garriga, L. & Ganley, I. G. Outstanding Questions in Mitophagy: What We Do and Do Not Know. J. Mol. Biol. 432, 206–230 (2020). 10.1016/j.jmb.2019.06.032

9 Martínez-Reyes, I. & Chandel, N. S. Mitochondrial TCA cycle metabolites control physiology and disease. Nat Commun 11, 102 (2020). 10.1038/s41467-019-13668-3

10 Allen, G. F., Toth, R., James, J. & Ganley, I. G. Loss of iron triggers PINK1/Parkin-independent mitophagy. EMBO Rep 14, 1127–1135 (2013). 10.1038/embor.2013.168

11 Lazarou, M. et al. The ubiquitin kinase PINK1 recruits autophagy receptors to induce mitophagy. Nature 524, 309–314 (2015). 10.1038/nature14893

12 Le Guerroué, F. et al. Autophagosomal Content Profiling Reveals an LC3C-Dependent Piecemeal Mitophagy Pathway. Mol. Cell 68, 786–796.e786 (2017). 10.1016/j.molcel.2017.10.029

13 Abudu, Y. P. et al. SAMM50 acts with p62 in piecemeal basal- and OXPHOS-induced mitophagy of SAM and MICOS components. J. Cell Biol. 220 (2021). 10.1083/jcb.202009092

14 Neuspiel, M. et al. Cargo-selected transport from the mitochondria to peroxisomes is mediated by vesicular carriers. Curr. Biol. 18, 102–108 (2008). 10.1016/j.cub.2007.12.038

15 Soubannier, V. et al. A vesicular transport pathway shuttles cargo from mitochondria to lysosomes. Curr. Biol. 22, 135–141 (2012). 10.1016/j.cub.2011.11.057

16 Prashar, A. et al. Lysosomes drive the piecemeal removal of mitochondrial inner membrane. Nature 632, 1110–1117 (2024). 10.1038/s41586-024-07835-w

17 Choong, C. J. et al. Alternative mitochondrial quality control mediated by extracellular release. Autophagy 17, 2962–2974 (2021). 10.1080/15548627.2020.1848130

18 Phinney, D. G. et al. Mesenchymal stem cells use extracellular vesicles to outsource mitophagy and shuttle microRNAs. Nat Commun 6, 8472 (2015). 10.1038/ncomms9472

19 Jiao, H. et al. Mitocytosis, a migrasome-mediated mitochondrial quality-control process. Cell 184, 2896–2910.e2813 (2021). 10.1016/j.cell.2021.04.027

20 Youle, R. J. & Narendra, D. P. Mechanisms of mitophagy. Nat. Rev. Mol. Cell Biol. 12, 9–14 (2011). 10.1038/nrm3028

21 Ding, W. X. & Yin, X. M. Mitophagy: mechanisms, pathophysiological roles, and analysis. Biol. Chem. 393, 547–564 (2012). 10.1515/hsz-2012-0119

22 Liberti, M. V. & Locasale, J. W. The Warburg Effect: How Does it Benefit Cancer Cells? Trends Biochem. Sci. 41, 211–218 (2016). 10.1016/j.tibs.2015.12.001

23 Else, T. et al. Adrenocortical carcinoma. Endocr. Rev. 35, 282–326 (2014). 10.1210/er.2013-1029

24 Miller, W. L. Steroid hormone synthesis in mitochondria. Mol Cell Endocrinol 379, 62–73 (2013). 10.1016/j.mce.2013.04.014

25 Munson, M. J. et al. GAK and PRKCD are positive regulators of PRKN-independent mitophagy. Nat Commun 12, 6101 (2021). 10.1038/s41467-021-26331-7

26 Wandinger-Ness, A. & Zerial, M. Rab proteins and the compartmentalization of the endosomal system. Cold Spring Harb. Perspect. Biol. 6, a022616 (2014). 10.1101/cshperspect.a022616

27 Baietti, M. F. et al. Syndecan–syntenin–ALIX regulates the biogenesis of exosomes. Nat. Cell Biol. 14, 677–685 (2012). 10.1038/ncb2502

28 Yim, W. W.-Y., Yamamoto, H. & Mizushima, N. A pulse-chasable reporter processing assay for mammalian autophagic flux with HaloTag. eLife 11, e78923 (2022). 10.7554/eLife.78923

29 Princely Abudu, Y., et al. NIPSNAP1 and NIPSNAP2 Act as "Eat Me" Signals for Mitophagy. Dev. Cell 49, 509–525.e512 (2019). 10.1016/j.devcel.2019.03.013

30 Hill, B. G. et al. Integration of cellular bioenergetics with mitochondrial quality control and autophagy. Biol. Chem. 393, 1485–1512 (2012). 10.1515/hsz-2012-0198

31 Tang, Z. et al. GEPIA: a web server for cancer and normal gene expression profiling and interactive analyses. Nucleic Acids Res. 45, W98–w102 (2017). 10.1093/nar/gkx247

32 Wang, T. & Rainey, W. E. Human adrenocortical carcinoma cell lines. Mol Cell Endocrinol 351, 58–65 (2012). 10.1016/j.mce.2011.08.041

33 Ishida, M., Arai, S. P., Ohbayashi, N. & Fukuda, M. The GTPase-deficient Rab27A(Q78L) mutant inhibits melanosome transport in melanocytes through trapping of Rab27A effector protein Slac2-a/melanophilin in their cytosol: development of a novel melanosome-targetinG tag. J. Biol. Chem. 289, 11059–11067 (2014). 10.1074/jbc.M114.552281

34 Yamano, K. et al. Endosomal Rab cycles regulate Parkin-mediated mitophagy. Elife 7 (2018). 10.7554/eLife.31326

35 Key, J. et al. Ubiquitylome profiling of Parkin-null brain reveals dysregulation of calcium homeostasis factors ATP1A2, Hippocalcin and GNA11, reflected by altered firing of noradrenergic neurons. Neurobiol. Dis. 127, 114–130 (2019). 10.1016/j.nbd.2019.02.008

36 Yim, W. W., Yamamoto, H. & Mizushima, N. A pulse-chasable reporter processing assay for mammalian autophagic flux with HaloTag. Elife 11 (2022). 10.7554/eLife.78923

37 Caserta, T. M., Smith, A. N., Gultice, A. D., Reedy, M. A. & Brown, T. L. Q-VD-OPh, a broad spectrum caspase inhibitor with potent antiapoptotic properties. Apoptosis 8, 345–352 (2003). 10.1023/a:1024116916932

38 TeSlaa, T. & Teitell, M. A. Techniques to monitor glycolysis. Methods Enzymol. 542, 91–114 (2014). 10.1016/b978-0-12-416618-9.00005-4

39 Cox, J. & Mann, M. MaxQuant enables high peptide identification rates, individualized p.p.b.-range mass accuracies and proteome-wide protein quantification. Nat. Biotechnol. 26, 1367–1372 (2008). 10.1038/nbt.1511

40 Tyanova, S. et al. The Perseus computational platform for comprehensive analysis of (prote)omics data. Nat. Methods 13, 731–740 (2016). 10.1038/nmeth.3901

41 Ritchie, M. E. et al. limma powers differential expression analyses for RNA-sequencing and microarray studies. Nucleic Acids Res. 43, e47 (2015). 10.1093/nar/gkv007

42 Dessau, R. B. & Pipper, C. B. [’’R"--project for statistical computing]. Ugeskr. Laeger 170, 328–330 (2008).

43 Kevin Blighe, S. R., Myles Lewis. EnhancedVolcano: publication-ready volcano plots with enhanced colouring and labeling, <https://github.com/kevinblighe/EnhancedVolcano> (2021).

44 Ge, S. X., Jung, D. & Yao, R. ShinyGO: a graphical gene-set enrichment tool for animals and plants. Bioinformatics 36, 2628–2629 (2020). 10.1093/bioinformatics/btz931

45 Colaprico, A. et al. TCGAbiolinks: an R/Bioconductor package for integrative analysis of TCGA data. Nucleic Acids Res. 44, e71 (2016). 10.1093/nar/gkv1507

46 Maglich, J. M., Kuhn, M., Chapin, R. E. & Pletcher, M. T. More than just hormones: H295R cells as predictors of reproductive toxicity. Reprod Toxicol 45, 77–86 (2014). 10.1016/j.reprotox.2013.12.009

47 Lin, C. W., Chang, Y. H. & Pu, H. F. Mitotane exhibits dual effects on steroidogenic enzymes gene transcription under basal and cAMP-stimulating microenvironments in NCI-H295 cells. Toxicology 298, 14–23 (2012). 10.1016/j.tox.2012.04.007

